# Histone H3 Orchestrates the Ubiquitination of Nucleosomal H2A by BRCA1/BARD1-UbcH5c Complex

**DOI:** 10.1101/2024.04.09.588726

**Authors:** Alexandra R. Goldman, Tejas Shah, Hedieh Torabifard

**Affiliations:** Department of Chemistry and Biochemistry, University of Texas at Dallas, 800 W. Campbell Rd., Richardson, TX

**Keywords:** Breast Cancer, DNA Damage, Histone Post-translational modification, Molecular Dynamics, Ubiquitin ligase

## Abstract

The Breast Cancer Associated Protein 1 (BRCA1) is a human tumor suppressor protein that commonly functions as ubiquitin ligase enzyme (E3) in the ubiquitination of the C-terminal H2A. BRCA1 enhances ubiquitin ligase activity by forming a heterodimeric complex with the BRCA1 Associated Ring Domain Protein (BARD1). The BRCA1/BARD1 complex works in concert with the ubiquitin-conjugating enzyme (UbcH5c or E2) to ubiquitinate one of the five lysines of the H2A C-terminal, ultimately promoting the repair of double-stranded DNA breaks. The mutations in the BRCA1-UbcH5c portion of the E3-E2 complex have been linked to breast and ovarian cancer. However, the mechanism of BRCA1/BARD1-UbcH5c complex ubiquitination at H2A is poorly understood, and the ubiquitination of exact lysine is debated. In this study, we sought to expand on the current research on H2A ubiquitination by using all-atom molecular dynamics simulations to model the BRCA1/BARD1-UbcH5c complex with the human ubiquitin protein (Ub). The Ub protein covalently bonds to the active site of E2, resulting in diminished flexibility of the E3-E2 complex with respect to the nucleosome core particle. The results of this study suggest a possible contribution of H3 in determining the preferred orientation of E2-Ub with respect to the H2A C-terminal lysines.

## Introduction

The nucleosome core particle (NCP) is a fundamental repeating subunit of chromatin found in the nucleus of eukaryotic cells.^[1]^ A single NCP consists of 147 base pairs of DNA wrapped around an octamer protein core consisting of two copies of H2A, H2B, H3, and H4 histone proteins.^[2]^ These histones contain a characteristic side chain, or tail, protruding out from the nucleosome with a high concentration of basic lysine and arginine residues. Although these tails do not affect the structure of NCP, they form the basis for histone post-translational modifications (PTMs) such as phosphorylation, acetylation, methylation, and ubiquitination.^[3,4]^ The PTMs affect chromatin structure in order to regulate biological processes such as gene expression, DNA repair, and chromosome condensation.^[1,2,4–8]^ As a consequence of the modification, the structural changes to the chromatin can act to include or even exclude certain protein complexes from interacting with DNA, thus altering its activity.^[5]^ The dysregulation of chromatin structure, such as that of H2A ubiquitination, has been associated with various cancers.^[2,3,6,7]^

The Breast Cancer Associated Protein 1 (BRCA1) is a human tumor suppressor protein that plays a vital role in DNA repair.^[9]^ Furthermore, BRCA1 functions as a ubiquitin ligase (E3), enhancing its E3 activity by forming a heterodimer complex with the BRCA1 Associated Ring Domain Protein 1 (BARD1).^[2,10,11]^ The structure of the heterodimer complex consists of four helix bundles containing four tetrahedrally-coordinated zinc metal centers.^[12]^ Defects in BRCA1/BARD1 have been found to be linked to familial breast and ovarian cancer and have therefore been an interest of study.^[10,12,13]^ In conjunction with UbcH5c, the ubiquitin-conjugating enzyme (E2), BRCA1/BARD1 can ubiquitinate specific lysine residues on the H2A C-terminal tail of the nucleosome.^[2,10,11]^ The structure of NCP with E3, E2, and ubiquitin (Ub) is shown in Figure 1. UbcH5c is a ubiquitin-conjugating enzyme found to function in concert with BRCA1 by covalently binding to it.^[11,14,15]^ Distortion of the BRCA1-UbcH5c complex significantly diminishes the E3 activity of BRCA1/BARD1 and, therefore, affects the structural integrity to ubiquitinate the H2A C-terminal tail.^[14]^

**Fig. 1:**
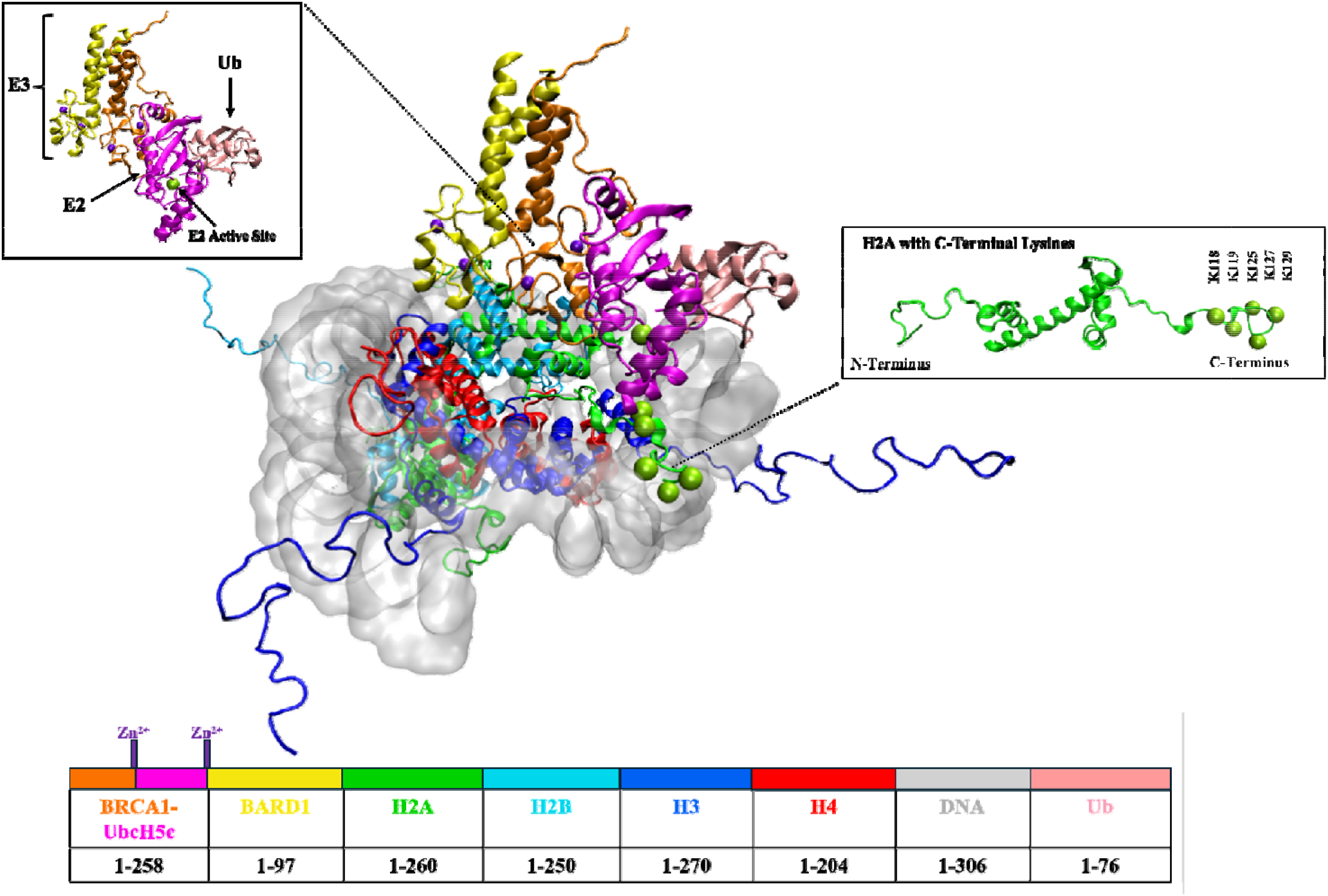
BRCA1/BARD1-UbcH5c bound to an NCP with H2A Histone. The figure depicts the H2A histone with the 5 prominent lysine residues susceptible to ubiquitination labeled and depicted as green spheres. BRCA1 (orange) and BARD1 (yellow) act in tangent as a ubiquitin ligase (E3) with UbcH5c (magenta), which is a ubiquitin-conjugating enzyme (E2). The structure also includes ubiquitin protein (pink) bonded to the active site of UbcH5c (K197). The table at the bottom of the figure contains the residue numbers present in each component of the structure. (Figure adapted from^[11]^)

Ubiquitination is a three-step process that attaches Ub to its desired substrate, as depicted in Figure S1. The process utilizes three enzymes: the ubiquitin-activating enzyme (E1), ubiquitin-conjugating enzyme (E2), and ubiquitin ligase (E3). The first step, activation, is an ATP-dependent process in which E1 catalyzes the formation of a thioester bond between the C-terminal carboxyl group of Ub and a cysteine on E1. In the next step, conjugation, Ub is transferred from E1 to E2, where Ub then bonds to the active site of E2. In the final step, ligation, E3 catalyzes the transfer of Ub from E2 to the target substrate, where the C-terminal glycine of Ub forms an isopeptide bond with a lysine on the target substrate.^[8,16]^

Studies have suggested that the BRCA1/BARD1-UbcH5c complex is sufficient for a preferential mono-ubiquitination of H2AK125, H2AK127, and H2AK129 at the disordered C-terminal H2A tail.^[7,10,12]^ In contrast, other ubiquitin ligases exist that display a different preference. For example, Ring1b/Bmi1 shows a preference for the mono-ubiquitination of H2AK118 and H2AK119.^[10,17]^ Our group has investigated the E3-E2 dynamics without Ub and proposed a mechanism by which the E3-E2 complex ubiquitinates the H2A C-terminal tail using all-atom molecular dynamic (AA-MD) simulations. The proposed mechanism identified a migration of H2AK127 and H2AK129 towards the active site of E2 when E2 moves away from the surface of the NCP, as a result of the increased flexibility in the H2A C-terminal tail. However, when E2 returns to the NCP surface, the mechanism proposed that H2AK118 and H2AK119 are then closer to the active site of E2.^[11]^ Lys residues on H2AK118 through H2AK129 are displayed on the depiction of the H2A histone tail in Figure 1. Considering the varying behavior from other E3 enzymes, significant research has been conducted on the ubiquitination preference of BRCA1/BARD1.^[7,10,11,12]^ However, due to the proximal positions of tail Lys residues and the dynamic nature of the disordered histone tail, it is difficult to characterize their conformational dynamics, as well as interactions with DNA and PTM factors.^[18]^ Given the function and regulations in the biological process, it is vital to understand the mechanism of ubiquitination of the H2A C-terminal tail by BRCA1/BARD1-UbcH5c.

In this study, AA-MD simulations have been used to investigate the influence of human Ub on the BRCA1/BARD1-UbcH5c/NCP dynamics and ubiquitination mechanism. The Ub was bonded to the active site of E2 (K197) *via* an isopeptide bond and compared to previous results to gain insight into the interaction of H2A with enzymes and Ub as well. It is important to note that while the active site of a UbcH5c is a cysteine, Witus *et. al* reported that the substitution of the E2 active site from a cysteine to a lysine stabilized the E3-E2/nucleosome interaction, allowing for more efficient purification of the structure.^[12]^ Hence, the active site of E2 was modeled as a lysine to remain consistent with the previous studies^[11]^ and the cryo-EM^[12]^ structure. In addition, the selectivity of ubiquitination by BRCA1/BARD1-UbcH5c in the presence of Ub has been characterized. Specifically, the impact and role of histone H3 in dictating ubiquitination have been assessed. Overall, this study intends to investigate the role of Ub on BRCA1/BARD1-UbcH5C dynamics and to enhance the understanding of ubiquitination machinery. Furthermore, the study of these proteins’ dynamics can be used to provide insight into the relationship between cancer incidence and Ub-dependent pathways and has promising applications in the field of therapeutic treatments. Indeed, MD simulations allow for a more thorough understanding of the mechanistic behavior of Ub-dependent pathways and, thus, allow for the potential of Ub-system targeted drugs as cancer therapeutics.^[19–23]^

## Methods

### Preparation of the Model

The structure used for this project was the structure of BRCA1/BARD1-UbcH5c bound to the NCP (PDB ID 7JZV^[12]^) that had been previously reported.^[11]^ The only change made to the structure was the addition of Ub while keeping other elements of the structure remained the same. Ub was attached to the active site of E2 via an isopeptide bond between the terminal glycine of Ub and the active site lysine (K197) of E2 (Figure S2). Based on the E2 active site substitution^[12]^, for this model, E2 and Ub form an isopeptide bond as opposed to the expected thioester bond.^[8,16]^

The structure of Ub was obtained from the cryo-EM structure of human tetraubiquitin (PDB ID 1F9J^[24]^). The monoubiquitin was prepared by separating the tetraubiquitin in VMD^[25]^, to obtain the coordinates of a single Ub protein. The protonation states of all titratable residues on Ub were determined by H++.^[26]^ The coordinates of Ub were then loaded with the E3-E2 NCP structure in VMD^[25]^, and the Ub was placed with the terminal glycine within range of the active site of E2. Using the AMBER Leap program^[27]^, Ub was bonded to the active site of E2 (K197) *via* an isopeptide using parameters obtained from Dr. Horn.^[28]^ The remainder of the system was kept consistent with the Ub-free system.^[11]^ The final structure has ∼420,000 atoms. Additionally, mutant systems were prepared in the same fashion as the WT, with the appropriate mutation of lysine residues to alanine. The detailed information for the model preparation can be found in the SI. Additionally, Table S1 contains the original residue index ranges for all components of the completed model.

### Simulation Details

The system was subjected to 2,000 steps of steepest decent minimization, followed by 3,000 steps of conjugate gradient minimization. In the first minimization phase, the protein and nucleosome were held fixed with a force constant of 500 kcal mol^-1^ Å^-1^. The second phase of energy minimization of the entire structure consisted of 20,000 steps. The system was then gradually heated for 1.5 ns to 300K under NVT conditions using a Langevin thermostat^[29]^ with a collision frequency of 1.0 ps^-1^. Finally, the system was equilibrated in NPT conditions using Berendsen weak coupling barostat^[30]^ for 2 ns with a weak restraint on the protein backbone before beginning production. The SHAKE algorithm^[31]^ was utilized for the purpose of constraining bonds containing hydrogen atoms. The Particle Mesh Ewald method was employed to accommodate long coulomb interactions^[32]^ and a 12 Å cutoff was applied for non-bonded interactions. Production snapshots were taken every 20 ps of the simulation in order to remain consistent with the previous work.^[11]^ All simulations were performed with the GPU version of pmemd in AMBER20.^[27]^ All visualization of the system was carried out using VMD.^[25]^ All analyses were performed using AMBER 20’s CPPTRAJ module^[33]^ or VMD.^[25]^ For salt bridge analysis, the residues were considered to form a salt bridge when the distance between the donor-acceptor atoms is < 3.2 Å and then the distance between the residues that formed said salt bridge as the simulation progress was monitored. The methodology of all analyses is consistent with that of the Ub-free system^[11]^ unless explicitly stated otherwise. Table S2 provides a summary of all simulations completed for this system, including the number of trials run and production time.

## Results and Discussion

### Stability and dynamics of the BRCA1/BARD1-UbcH5c/NCP with bound ubiquitin

The overall structural stability of the WT system was determined by calculating the RMSD of the backbone for BRCA1/BARD1-UbcH5c, Ub, DNA, and the 8 core histones (with and without tails). The results of RMSD are depicted in Figure S3 and suggest that the system was well equilibrated. The RMSDs of the histone cores with tails are significantly larger due to the large fluctuations of flexible disordered tails. Notably, the RMSD of H3 with tails is approximately two to four times larger than the RMSDs of H2A (Fig. S4a-c) across all WT trials. The RMSD for the BRCA1/BARD1-UbcH5c complex is smaller (< 5 Å) than the RMSD reported for the BRCA1/BARD1-UbcH5c complex in the Ub-free system.^[11]^ This, therefore, indicates that the presence of ubiquitin constrains the E3-E2 dynamics, resulting in less structural flexibility.

The RMSF of the sidechains of BRCA1-UbcH5c and BARD1 (shown in Fig. 2a,b) were found to follow a similar fluctuation pattern to those observed for the Ub-free system.^[11]^ The disordered terminal residues of the proteins observed the highest fluctuations, suggesting the flexibility of these tail residues. Interestingly, the RMSF of the sidechains of Ub (Fig. 2c) exhibits a greater fluctuation of interior residues’ side chains compared to the other protein components of the system. The residues of Ub that exhibit the highest fluctuations (RMSF > 5 Å) are found on the surface of Ub, interacting with the H3 N terminus tail, and are shown in cyan in Figure 2d. Furthermore, Figure S5a shows that the RMSF of H3 for the first 20 residues of the protein exhibits a large fluctuation (RMSF > 5 Å). Interestingly, this region of high fluctuation on H3 is the portion of the N terminus of the disordered tail that is seen to interact with Ub or the NC in the simulations. Figure S5b shows that the residues located at the C-terminal of H2A also exhibit a large fluctuation (RMSF > 5 Å), which is expected due to the flexible nature of histone tails.

**Fig. 2:**
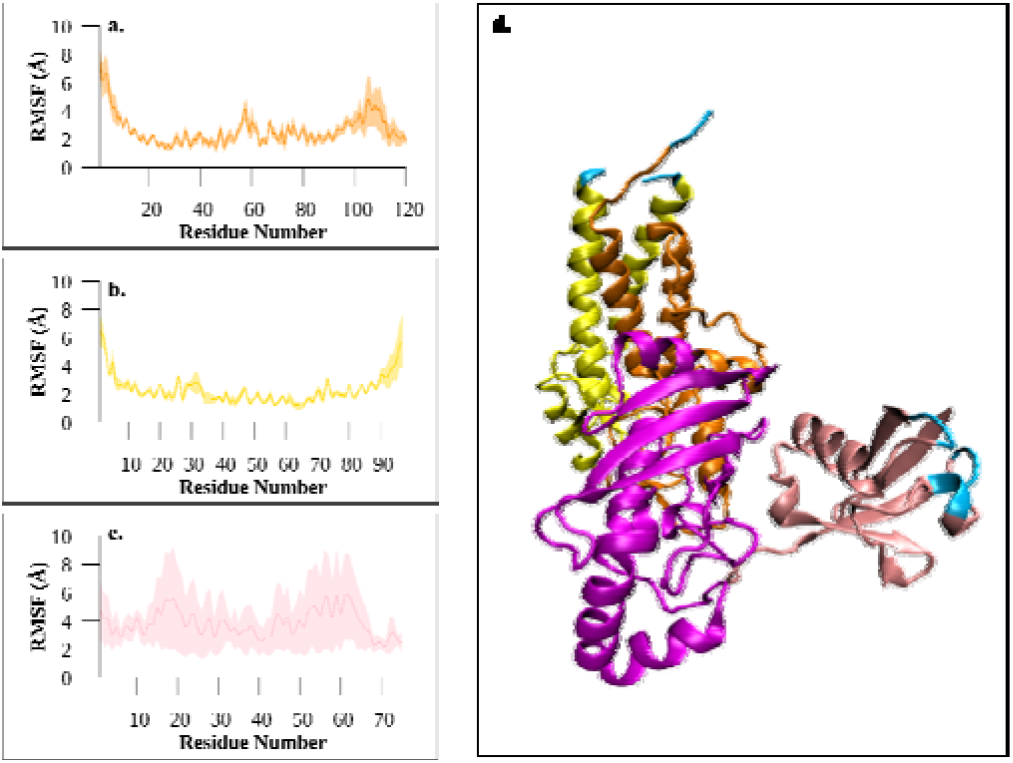
Fluctuations of the System. RMSF of (a) BRCA1/UbcH5c, (b) BARD1, (c) Ub for WT simulations. (d) BRCA1/BARD1/UbcH5c-Ub complex with cyan indicating regions of large fluctuation (RMSF > 5 Å). All shading of plots indicates the standard deviation of the data set.

### Tilt angle and distance analysis reveal the movement of the BRCA1/BARD1-UbcH5c/Ub

The tilt angle of BRCA1 and BARD1 relative to H2B in the cryo-EM structure (in which Ub was not present) was found to be approximately between 84° and 86°.^[11,12]^ To evaluate the influence of Ub on the movement of BRCA1/BARD1, we calculated the tilt angle for the systems. The angle is defined as the angle formed between a vector passing through an alpha helix of H2B and an alpha helix on BRCA1 or BARD1 to form the BRCA1 tilt angle and BARD1 tilt angle, respectively.^[11]^ A larger tilt angle would suggest that the heterodimer is tilted towards the NCP surface containing the H2A C-terminal tail. In contrast, a smaller tilt angle implies the heterodimer is tilted farther from the H2A C-terminal tail, as visualized in Figure 3a. The tilt angles shown in Figure 3b show that the tilt angle for BRCA1 is distributed around 90°, while BARD1 is distributed around 95°. The tilt angles for BRCA1, BARD1 for individual trials, and BRCA1/BARD1 tilt angles as the progression of simulation time are shown in Figures S6-S8. The larger tilt angles were consistently observed in all the trials of the Ub bound system, implying that the presence of Ub plays a role in orienting or “pulling” the heterodimer towards the NCP surface. Furthermore, the tilt in each case shows a narrow distribution in a span of 30° for each tilt compared to our previous results with the Ub-free system, which showed a range of 100°.^[11]^ In this instance, the constricted large tilt angle may suggest that Ub not only orients the E3-E2 complex but also constrains E3-E2’s previously defined “hinging” motion^[11]^, resulting in a closer orientation of the complex to the NCP surface. The different motions of E3-E2, including hinging motion, will be discussed in more detail in a later section.

**Fig. 3.**
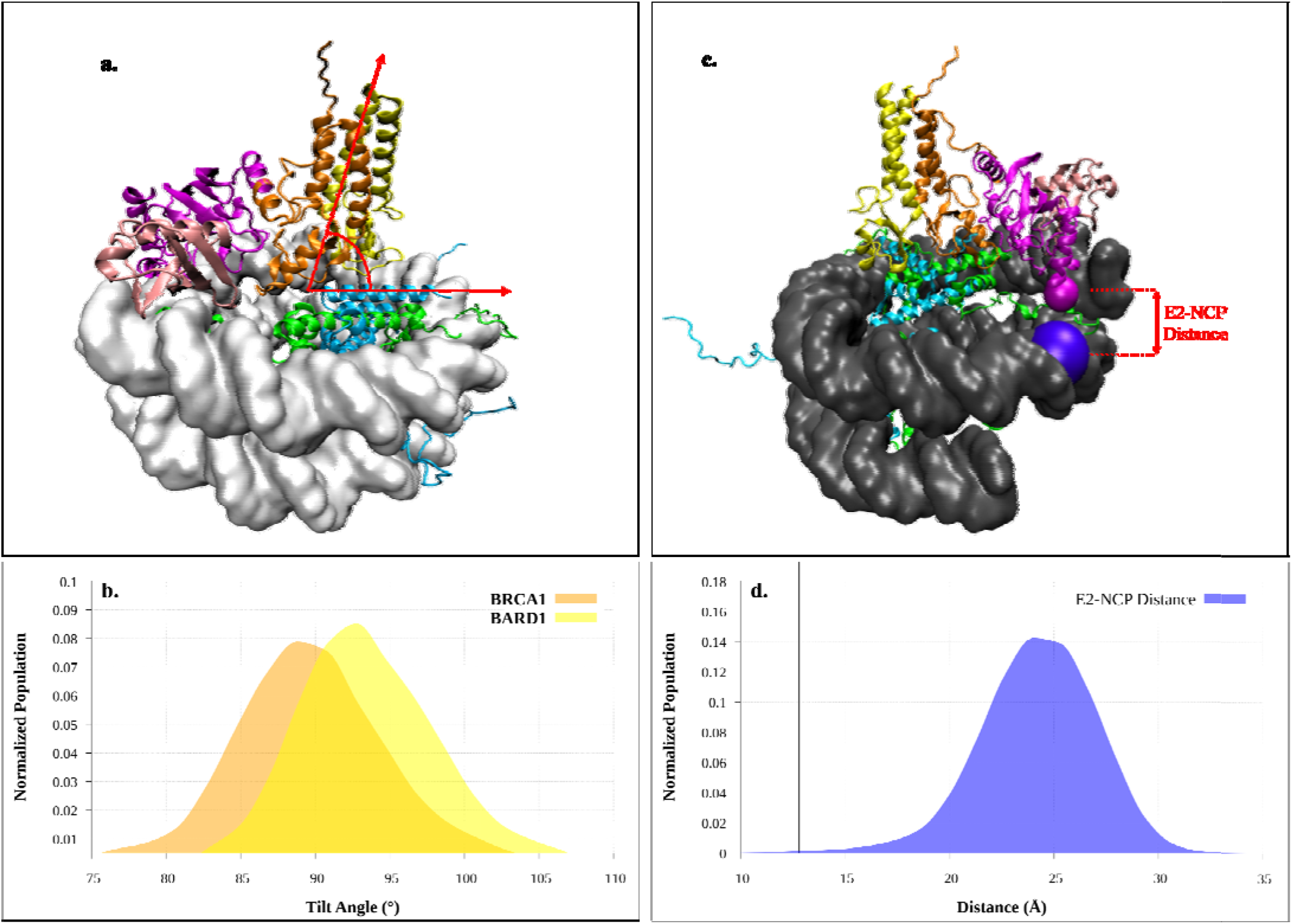
BRCA1/BARD1 Tilt Angles and E2-NCP Distance. (a) The tilt angle is defined by a vector passing through an alpha helix of H2B (cyan) and an alpha helix passing through BRCA1 (orange) or BARD1 (yellow). A larger tilt angle suggests the heterodimer is tilted towards E2 (magenta) and Ub (pink). H2A (gree) is also included in the diagram. (b) The normalized tilt angle of BRCA1 (orange) and BARD1 (yellow) across all WT trials. (c) E2-NCP distance is defined as the distance between the alpha carbon of D242 of E2 (magenta) and the P of residue 1443 on DNA (purple). (d) The normalized E2-NCP distance across all WT trials reported in Å. The black line indicates the starting distance of 12.80 Å as reported of the cryo-EM structure.^[11,12]^

To obtain further insight into the behavior of E2 with respect to the NCP surface, we performed distance analysis. The initial distance, as reported in the cryo-EM structure, is 12.8 Å, and the distance analysis of the Ub-free system shows a bimodal curve with normalized distances of ∼20 Å and 40 Å.^[11]^ However, in the Ub-bound system, the E2-NCP distance increases to 35 Å, normalizing around 25 Å. Figure 3c provides a visualization of the E2-NCP distance, and Figure 3d shows the E2-NCP distance for the Ub-bound system, which is consistent with the lower range reported for the Ub-free system.^[11]^ The lack of the bimodal curve in the Ub-bound system suggests that the presence of Ub limits the flexibility of E2, coercing E2 to remain closer to the NCP surface for the entirety of the simulations. This observation is consistent with the results of the RMSD, and tilt angle analysis discussed previously. As depicted in the diagram (Fig. 3c), when the E2-NCP distance is smaller, E2 is oriented closer to the NCP surface. The shorter E2-NCP distance (shown in Fig. 3d) complements the results of the tilt angle analysis in suggesting that Ub limits the flexibility of the E3-E2 complex, such that it is closer to the NCP and, therefore, closer to the C-terminal tail of H2A.

### H3-Ub interaction presents a possible modulation of BRCA1/BARD1-UbcH5c dynamics

Across the three WT trials, there is a critical difference in the behavior of the H3 histone N-terminal tail. In 33% of the simulation time, the first 20 residues of the H3 N-terminal tail stretch upward and interact with the Ub protein. This interaction between the H3 tail and Ub was established within the first 100 ns of the first WT trial. The interaction is further supported by the salt bridge analysis (Fig. 9a), in which the N-terminal tail of H3 forms salt bridges with 7 different residues on Ub (E16, E18, D21, E24, E51, D52, and D58) and H3 settles within <40 Å of these residues after the salt bridge was formed. In contrast, for the remaining 67% of the simulations, the H3 tail only interacts with the NCP and not with the Ub protein. Furthermore, the salt bridge analysis suggests that when H3 does not interact with Ub, it interacts with limited residues on E2 (Fig. S9b, c). Additionally, in the second WT trial (Fig. S9b), H3 forms a salt bridge with H2AE121 on the H2A C-terminal tail, in which the residues remain relatively close (<20 Å) for the duration of the trial. This salt bridge suggests that in the case of WT trial 2, H3 is closer to the protein core, specifically H2A. Depictions of possible H3 interaction (for each of the three trials) are found in Figure 4 a-c. Additionally H3 contact frequency (Fig. S10) shows that when the H3-Ub interaction is absent, contact between H3 and Ub is significantly diminished (Fig. S10a). The contact frequency: however, reveals that in the second WT trial, H3 is indeed within proximity to H2A, but not in the third WT trial (Fig.S10b). It is important to note that in half of the H3-NCP interaction simulation times in WT, the H3 N-terminal tail reaches upward and attempts to interact with Ub; however, it is unsuccessful and eventually interacts with the outer portion of DNA on the NCP instead (Fig. 4c). This contrasts with the H3-NCP interaction in trial 2 (Fig. 4b) as H3 has more contact with the protein core (the H2AE121-H3K14 salt bridge). This would suggest that when H3 is unsuccessful in interacting with Ub, its proximity to the NCP’s protein core diminishes. The contact of H3 with the areas of interest in the protein (Ub and H2A) over all WT trials can be found in Table S3. The table summarizes that the H3-Ub contact dominates in the first WT trial, while in the second trial, the H3-NCP contact dominates. As mentioned previously, H3 is unable to contact with Ub and NCP in trial 3, as the tail becomes localized on the exterior of the DNA (Fig. 4c). Furthermore, the salt bridge analysis supports the diminished contact of H3 in trial 3, as the H3 N-terminal tail is only able to form 2 salt bridges with E2 and Ub (Fig. S9b). Specifically, the salt bridge that forms between H3 and Ub in the third trial shows the largest change in distance between the two salt bridges formed, as H3 eventually moves away from Ub, and is caught in DNA. Furthermore, the H3 contact frequency (Fig. S10) supports that H3 has diminished contact with both Ub and H2A in the third WT trial. The distal impact of histone H3 on BRCA1/BARD1 mediated ubiquitination has also been reported by Witus *et al*. ^[12,17,34]^ However, unlike our results, their observations implied the involvement of the ordered region of histone H3.

**Fig. 4.**
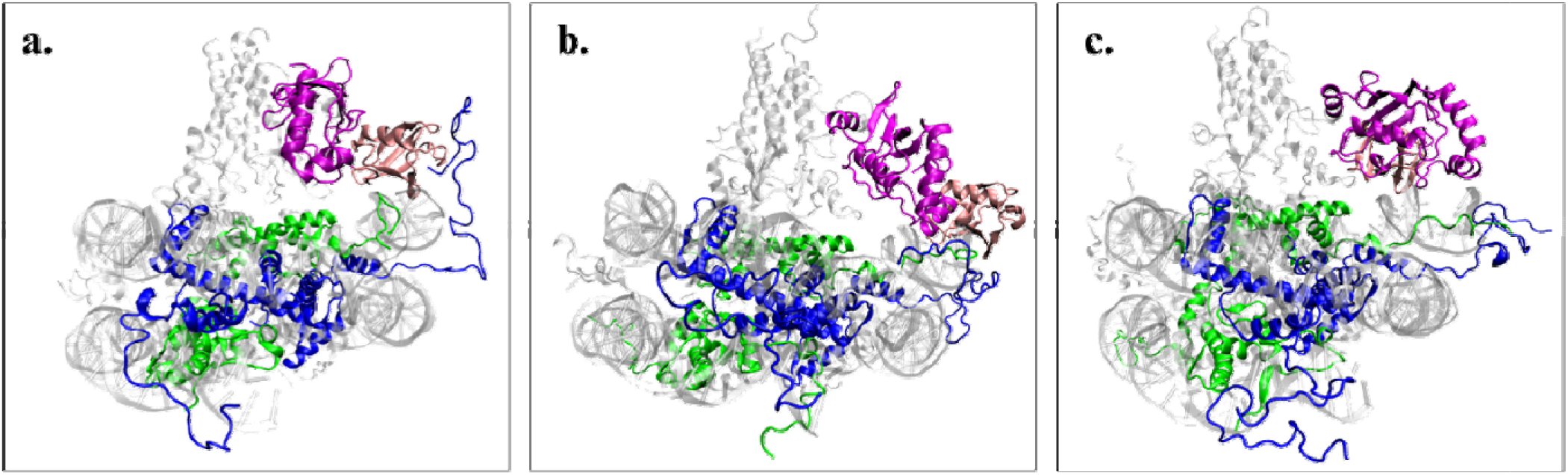
Behavior of WT trials at 100 ns. (a) WT-trial 1: H3 (blue) interacts with Ub (pink), causing an upward tilt of UbcH5c (magenta). (b) WT-trial 2: H3 interacts with the NCP (gray) rather than Ub. (c) WT-trial 3:H3 interacts with the NCP. H2A is shown in green.

The effect of the H3-Ub interaction impacts the tilt angle of both BRCA1 and BARD1 over the duration of their respective simulations (Fig. S6-S8). Across the three trials, the tilt angle for both BRCA1 and BARD1 is found to be distributed at the largest angle for 33% of the simulation time with an H3-Ub interaction. In contrast, the remaining 67% of the simulation time with no H3-Ub interaction depicts a lesser angle in comparison. The tilt angle over time plots (Fig. S8) shows that the tilt’s behavior varies depending on the behavior of H3. In the first trial (Fig. S8a), the tilt angle of both BRCA1 and BARD1 increases and eventually stabilizes at about 90° for both BRCA1 and BARD1, suggesting that the H3-Ub interaction brings the heterodimer closer to the C-terminal of H2A during the progression of the interaction. Inversely, in the third trial (Fig. S8c), the tilt angle decreases from its starting position and stabilizes at an angle of about 70° for BRCA1 and 85° for BARD1, making the heterodimer further from the H2A tail than in trial 1. In all trials, the BRCA1 tilt always starts at a lower degree than its BARD1 counterpart; however, when H3 immediately interacts with the NCP (WT trial 2), the BRCA1 tilt angle increases while the BARD1 tilt decreases, and both stabilize at an angle of 85° (Fig. S8b). The varying behaviors of both BRCA1 and BARD1 tilt angles across the three trials suggest that the movement of the heterodimer with respect to the NCP is reliant on whether H3 is able to interact with Ub or not. The normalized tilt angles of individual trials (Fig.S6-S7) further support that when the H3-Ub interaction is present (Fig. S6a and S7a), the tilt angle is larger compared to the other trials, suggesting that H3 plays a role in bringing the heterodimer, and thus Ub, closer to the surface of the NCP containing the H2A C-terminal tail. This trend in tilt angle suggests that H3 may play a role in dictating the orientation of BRCA1/BARD1 with respect to H2A.

### Asymmetric dynamics of E2 and Ub with respect to the NCP

To compare the conformational changes of the Ub-bound system to the Ub-free system^[11]^, Principal Component Analysis (PCA) was performed to define large-scale conformational changes in the system. The PCA for this system was performed in the same manner as the Ub-free system, and PCs accounting for > 10% of the system conformations were examined. For the purpose of consistency, movements of the system will be referred to as “tilting, sliding, hinging and rotational” modes as defined in the previous publication.^[11]^ The distinction was the PCA was performed on the BRCA1/BARD1-UbcH5c/Ub portion of the system with respect to the NCP, and the PCA was conducted on all WT simulations. Table S4 provides information on the principal modes present. In Table S4, mode A correlates to the “seesaw” motion of UbcH5c/Ub, which accounts for the largest contribution of movement across all simulations. Mode B is defined as the inverse rotation movement of UbcH5c and Ub about the BRCA1/BARD1 on the NCP surface. Finally, mode C represents the upward movement of E2 with a sliding Ub present in the first trial of the simulations. Figure 5a provides visualizations of all modes. The movement of the BRCA1/BARD1 heterodimer across all WT systems exhibits the same “tilting” motion as observed in our previous studies.^[11]^ However, the intensity of the tilt varies among the simulations. For the PCs present in the highest variance, as depicted in Table S4, the tilting motion of the heterodimer is most prominent for the second trial, where H3 almost immediately interacts with the NCP. This implies that when H3 interacts with the NCP, the heterodimer returns to a smaller tilt angle rather than remaining at a higher tilt (Fig. S8b). For the remaining simulations, H3 does not immediately interact with the NCP; thus, a different tilt behavior is observed (Fig. S8a, c). When the H3-Ub interaction occurs, the heterodimer remained at a higher tilt angle for the majority of the simulation (Fig. S6a, S7a). This would confirm the less prominent tilt seen in the PC of trial 1, as it does not drastically shift from the starting tilt. Furthermore, when H3 comes close to Ub to interact, the heterodimer remains at the higher tilt angle longer than that observed in the portion of the simulations with no H3-Ub interaction (Fig. S8a, c). Since Ub is bonded to UbcH5c’s active site, the movements of both UbcH5c and Ub are grouped together. All significant modes observed for 33% of the simulation time with the H3-Ub interaction exhibit a sliding motion of Ub, similar to that exhibited by E2 in mode 2 of the Ub-free system.^[11]^ Interestingly, for the majority of modes present for this first WT trial, E2 hinges towards or away from the NCP surface, causing the subsequent sliding of Ub up or down in the reverse direction of E2. In the other 66% of the simulation time, a “seesaw” like motion is observed between E2 and Ub for a majority of those simulations’ variances. For this “seesaw” motion, E2 moves down to the NCP surface while Ub moves up and vice versa. The up and-down movement of E2 is similar to the hinging motion defined previously.^[11]^ The second major PC in the first two trials of the simulations consists of the rotation mode in E2 previously observed.^[11]^ When E2 rotates about the heterodimer on the NCP surface, it seems similar to the inverse behavior of the “seesaw” motion, causing Ub to rotate about the heterodimer in the opposite direction of E2.

**Fig. 5.**
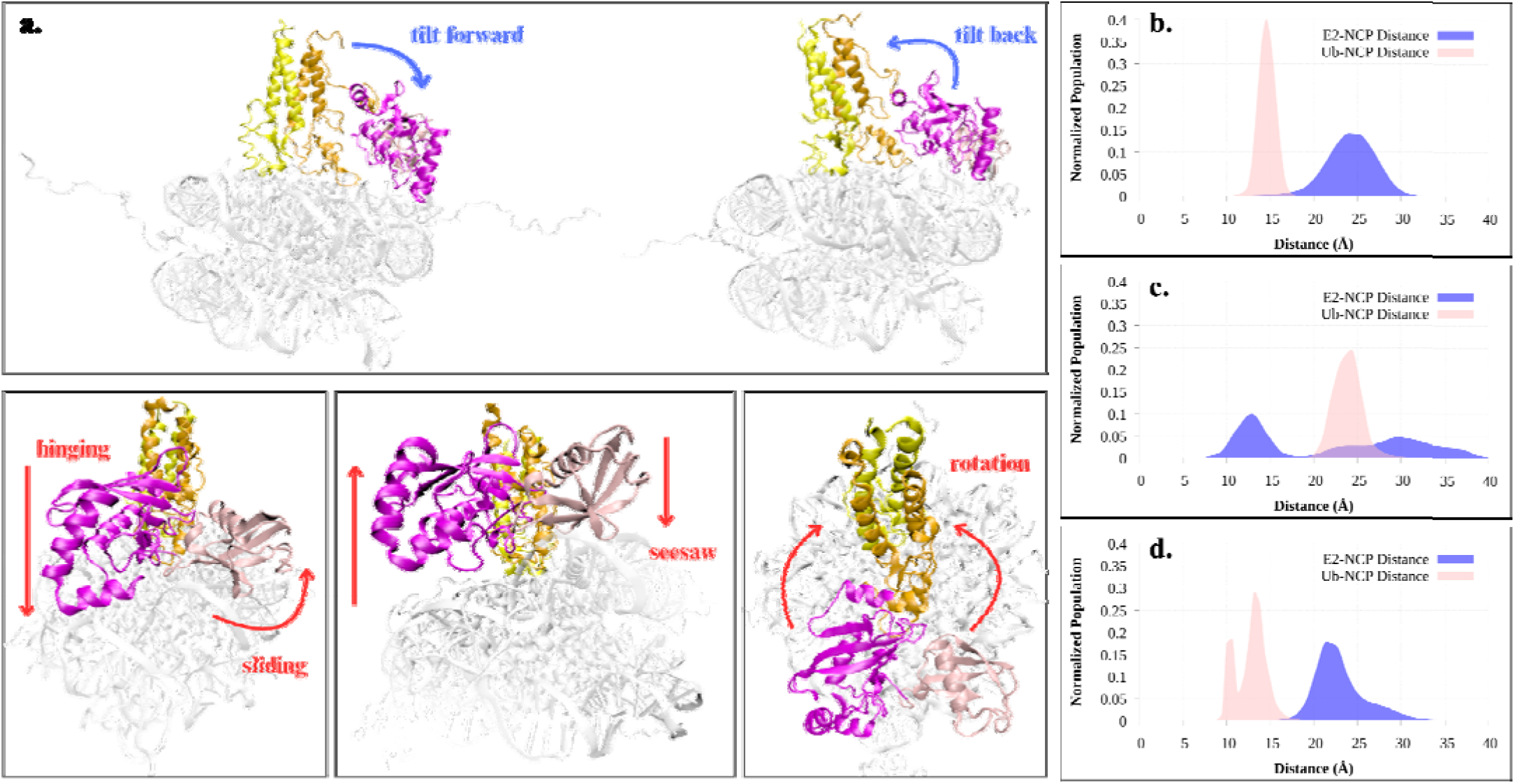
Asymmetric Movements of E2 and Ub. (a) Visual summary of all PCA modes including tilting, hinging, sliding, rotating and newly defined “seesaw” of RCA1/BARD1-UbcH5c/Ub in WT. E2-NCP (blue) and Ub-NCP (pink) distance of (b) WT trial 1. (c) WT trial 2. (d) WT trial 3. The Ub-NCP distance is defined as the distance between the alpha carbon of A46 on Ub and the phosphate on residue 1651 of DNA. The starting Ub-NCP distance was 11.55 Å.

To better compare the movement of E2 and Ub with respect to the NCP, a Ub-NCP distance analysis was conducted in the same methodology as the E2-NCP distance. The E2-NCP distance of individual WT trials (Fig. 5b-d) indicates that in the presence of the H3-Ub interaction, E2 normalizes at a larger distance from the NCP surface that contains H2A (Fig. 5b) compared to the trials with no H3-Ub interaction (Fig. 5c, d). Interestingly, the reverse is true for the Ub-NCP distance. When the H3-Ub interaction occurs, the distance of Ub to the NCP surface with H2A is less than that of E2 but is still larger than the starting distance of Ub. (Fig. 5b). In contrast, when the E2-NCP distance is at its shortest (Fig. 5c), the Ub-NCP distance is at its longest. The third WT trial in which H3 attempts to interact with Ub but is unsuccessful (Fig.4c) shows an interesting behavior. While H3 eventually interacts with the NCP in the third trial, BRCA1 remains tilted away from the H2A C-terminal (at a smaller tilt angle), and E2 remains farther (Fig. 5d) compared to trial 2 (Fig. 5c). This larger E2-NCP distance for the third trial then results in a relatively smaller Ub-NCP distance (Fig. 5d). In contrast, the H3-Ub interaction (trial 1) promotes a sustained larger BRCA1 tilt angle (Fig. S6a), and a constrained Ub-NCP distance (Fig. 5b). Therefore, the failure of the H3-Ub interaction to occur in the third trial is substantiated by the smaller BRCA1 tilt angle, and wide distribution of the Ub-NCP distance (∼10 Å) (Fig. 5d). Since the dynamics of the Ub-free system predicted that when E2 moves away from the NCP, the H2A C-terminal tail become more flexible and allows for the migration of H2AK127 and H2AK129 to the E2 active site^[11]^, the H3-Ub interaction may play a role in the ubiquitination process. The H3-Ub interaction results in the movement of E2 away from the NCP and Ub towards the NCP, and thus, may promote the movement of the last three lysine to the active site of E2. Furthermore, the inverse movement of E2 and Ub suggest that when E2 moves away from the NCP, Ub moves closer to the NCP, and thus, H2A.

The inverse motion of Ub with respect to E2 can therefore be corroborated by the inverse relationship between the E2-NCP and Ub-NCP distances (Fig. 5b-d). The prominence of the newly defined “seesaw” motion is representative of the inverse distance trends of E2 and Ub with respect to NCP, and further shows that if E2 moves upward, Ub moves down, and vice versa. The inverse motions of the two proteins may be indicative of that when E2 moves away from the NCP, Ub is able to be oriented closer to the NCP for its eventual transfer to H2A. In summary, the modes present for the Ub-bound system are similar to those present in the Ub-free system, however; the presence of Ub reflects the constraints on the dynamics of E2, and the dependence of Ub’s movements on E2 for the preparation of the Ub transfer.

### Interactions between UbcH5c (E2) and the H2A C-terminal lysines

In order to investigate the interactions between E2 and the H2A C-terminal lysines, the contact frequency between the H2A C-terminal lysines and residues on UbcH5c was determined. The contact frequencies reported for individual trials can be found in Figure S11. When comparing the contact frequencies across the WT simulations, the 33% of the simulation time when the H3-Ub interaction exists, H2AK125 favors interaction with E2, whereas the rest of the WT simulations show that H2AK119 and H2AK118 favor interactions with E2. The findings of contact frequency were further corroborated by a distance analysis between the C-terminal H2A lysines and the E2 active site, which revealed a similar preference for H2AK125 and H2AK119 depicted in Figure 6. The preference for H2AK125 exists in the presence of the H3-Ub interaction, while the preference for H2AK119 occurs in the absence of the H3-Ub interaction. Moreover, in the presence of H3-Ub interaction, H2AK127 and H2AK129 are in a closer proximity to the E2 active site compared to the other WT trials (Fig. 6a). The ubiquitin transfer to the terminal three lysines; H2AK125, H2AK127, and H2AK129 by BRCA1/BARD1-UbcH5c complex has been reported in the literature.^[7,10,12,17,34]^ Therefore, the selectivity for H2AK125 in the presence of the H3-Ub interaction could be representative of the results reported by experimental findings.

**Fig. 6:**
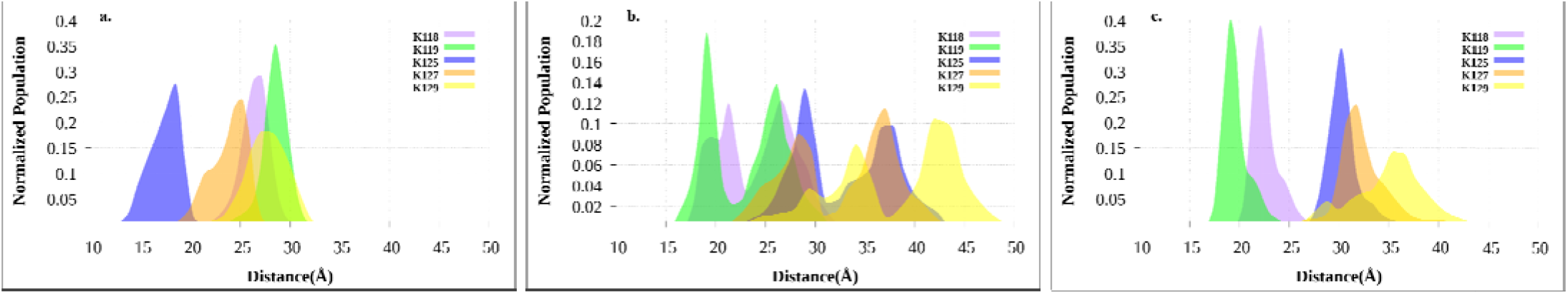
Distance Between C Terminal Lysines and E2 Active Site Across WT Trials. Distance between K118 (purple), K119 (green), K125 (blue), K127 (orange) and K129 (yellow) with the E2 active (K197) for (a) WT trial 1(b) WT trial 2 (c) WT trial 3. The distance is the distance of the center of mass of lysine residue with respect to the E2 active site center of mass.

Additionally, in 67% of the WT simulation time in which there is no H3-Ub interaction, residues E2E234 through E2K245 exhibit significant contact frequencies with H2AK125, H2AK127, and H2AK129 (Fig. S11b,c). However, Figure S11c, which corresponds to the WT trial 3 with the largest E2-NCP distance in the absence of the H3-Ub interaction (Fig. 5d), shows a larger contact frequency percentage with the three terminal lysines compared to contact frequencies obtained for WT trial 2 (with no H3-Ub interaction) shown in Figure S11b.

This may then suggest that the terminal 3 lysines on H2A are not entirely dependent on the H3-Ub interaction to approach the E active site; however, the absence of the H3-Ub interaction still appears to limit the flexibility of these lysines compared to the dynamics when the H3-Ub interaction is present (Fig. 6). The presence of the H3-Ub interaction may then suggest that the interaction between H3 and Ub plays a role in altering the selectivity of the ubiquitination of the H2A C-terminal. This possibly explains why H2AK125, which beh ves as an anchor in the absence of Ub^[11]^, seems to be favored for ubiquitination when the H3-Ub interaction occurs. In fact, these findings may also support the discrepancies observed for Ub-conjugation at H2A lysines reported during in vitro and cellular assays.^[7,12]^ Furthermore, it is critical to recall that H3 itself can undergo PTMs; for example, the acetylation of H3 can stimulate the de-ubiquitination of a monoubiquitinated H2A.^[35]^ In addition to the acetylation of H3, researchers have reported that H3 can undergo mono-ubiquitination mediated by the E3/E2 hybrid enzyme, UBE2O.^[36]^ In summary, the PTMs of H3 may play a role in the desire of H3 to interact with Ub.

To further quantify the interactions between H2A and E2 and to better understand the dynamics of H2A, a salt bridge analysis was once again conducted. For this analysis, salt bridges formed with residues E2D228 and E2D229 of the UbcH5c loop region were investigated. The interest in E2D228 and E2D229 arises from their strong interactions with the active site of E2 (K197) and the H2A C-terminal lysines observed in our previous report.^[11]^ Since Ub is bound to the active site of E2, the interactions between E2D228 and E2D229 with E2K197 were expected to be diminished; however, their behavior was consistent with that of contact frequency results. Results from the salt bridge analysis for all three WT trials can be found in Figure S12a-f. In simulations with H3-Ub interaction, H2AK125 is the only C-terminal Lys that forms a salt bridge with both E2D228 and E2D229 (Fig. S12a,d). For the remainder of the simulation after forming the salt bridge, H2AK125 stays within 25 Å of E2D228 and 15 Å within E2D229. Interestingly, this salt bridge was not detected during the other simulations, suggesting that H2AK125 did not exhibit the flexibility required to approach the E2 active site. Alternatively, salt bridges between E2D228 and E2D229 with both H2AK118 and H2AK119 were established in the absence of the H3-Ub interaction (Fig. S12b,c,e,f). Additionally, the distance of H2AK119 to E2D228 and E2D229 was shorter (5 to 10 Å) than compared to their di tances with H2AK118 over the entirety of the simulations following the formation of the salt bridge. The results of the salt bridge analysis suggest that in the absence of the H3-Ub interaction, the flexibility of the C-terminal H2A lysines is limited such that only H2AK118 and H AK119 have the ability to approach the E2 active site. Therefore, we propose that the occurrence of the H3-Ub interaction affects the flexibility of the H2A C-terminal tail, and ultimately the lysine preference for ubiquitination.

### Mutants provide insight into the role of the H3-Ub interaction

In order to further investigate the dynamics of the system, we prepared a double mutant (H2AK118A and H2AK119A) and a triple mutant (H2AK125A, H2AK127A, and H2AK129A). The mutations for the multiple mutants were grouped such that the dynamics could be examined when mutations were present in the ordered (double mutant) and disordered (triple mutant) regions of the H2A C-terminal tail.

The results of tilt angle and E2-NCP distance for these mutants are depicted in Figure 7 a-c. The double mutant has a slightl larger BRCA1 tilt compared to the WT (Fig. 7a); however, this is not the case for the BARD1 tilt. Notably, the BARD1 tilt for the double mutant system is much lower (∼20°) than its BRCA1 counterpart for the double mutant. Additionally, the BARD1 tilt for the double mutant is ∼10° lower than the WT (Fig. 7b). This may suggest that there is an asymmetric movement of the heterodimer to orient the BRCA1 towards the NCP in the double mutant. Additionally, the triple mutant exhibits a substantially larger E2-NCP distance than the double mutant (by ∼10-15 Å) (Fig.7c). The contact frequency between E2 and H2A for the multiple mutant systems supplements this discovery (Fig. S13a,b). In the case of the double mutant system, contact frequencies are quite large (> 40%) compared to the WT for the H2A tail with E2 (Fig. S11). In contrast, the contact frequencies observed for the triple mutant system are significantly diminished when compared to the WT. Furthermore, when these mutants were visualized, as depicted in Figures 7 d and e, the fate of the H3 histone tail varies. In the case of the double mutant, the H3 tail remains close to the Ub/NCP for the duration of the simulation (Fig. 7d). On the other hand, the H3 tail in the triple mutant remains extended outward from the NCP for the entirety of the simulation and does not interact with the protein (Fig. 7e).

**Fig. 7.**
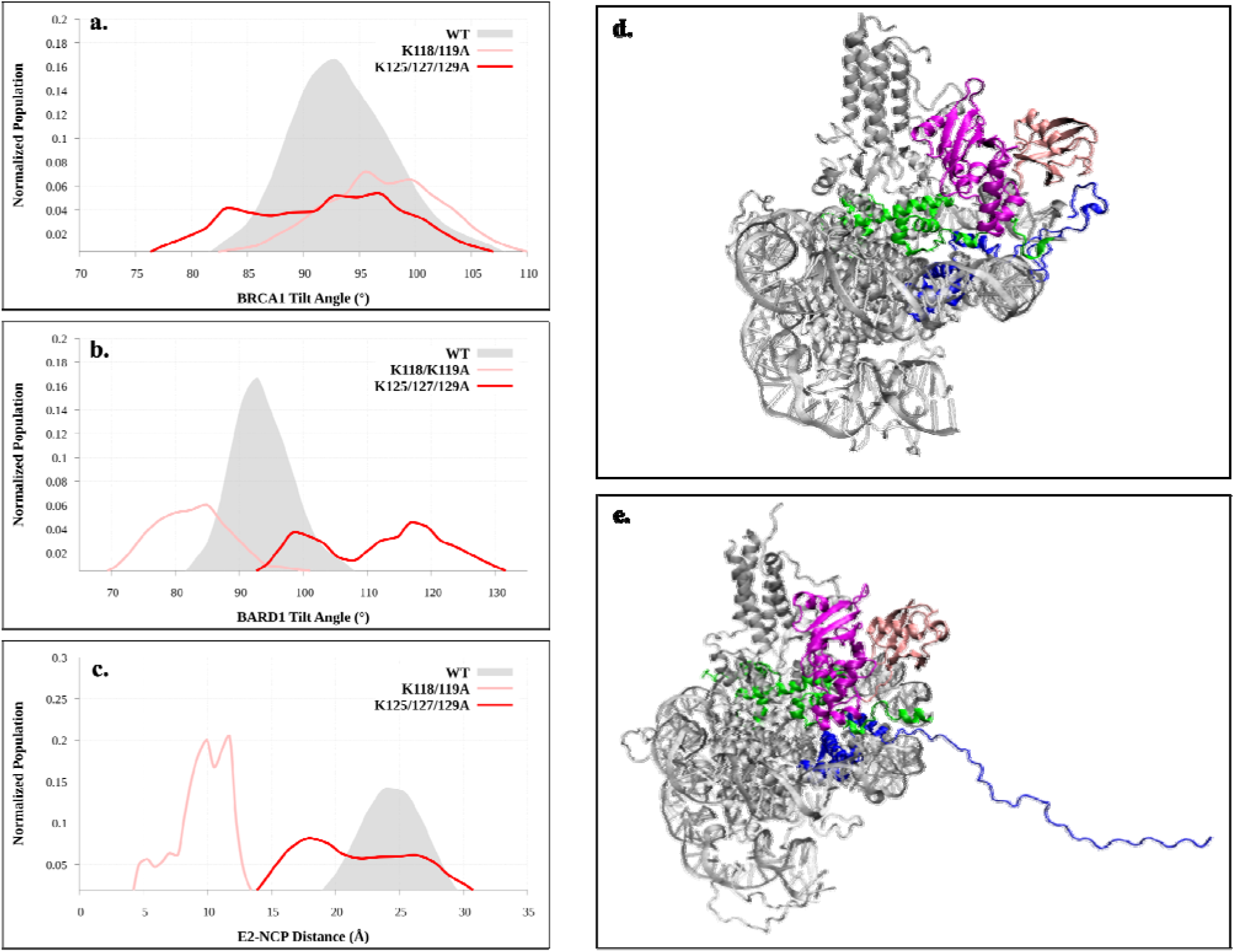
Results of tilt Angle and E2-NCP Distance for Double and Triple Mutant Systems. (a). The WT BRCA1 tilt angle (gray) versus that of the double mutant (pink) and the triple mutant (red). (b) The WT BARD1 tilt angle versus the double and triple mutant systems. (c) The E2-NCP distance of WT versus double and triple mutant systems. (d) The double mutant system at t=100 ns observes the H3 histone tail oriented towards Ub and the NCP, whereas (e) the H3 tail of the triple mutant at t=100 ns is away from the NCP.

In order to gain further insight into the role of H3, H3 contact frequency with both H2A and Ub was investigated. Figures S14 a and b show the contact frequency between the H3 N-terminal tail and the C-terminal H2A lysines (Fig. S14a) and areas of high fluctuation of Ub (Fig. S14b). The contact frequency results of double mutant systems display the higher contact of the H3 N-terminal tail with both H2A (Fig. S14a) and Ub (Fig.S14b) compared to the triple mutant (which has a H3-Ub and H3-H2A contact frequency of 0%). While the double mutant undergoes both the H3-Ub and H3-H2A contact for the duration of its simulation time, the H3-Ub contact dominates (by ∼10%) as shown in Table S3. It is crucial to recall that in the presence of the H3-Ub interaction, E2 moves away from the NCP and Ub moves towards the NCP, allowing for greater flexibility of the last three lysines on H2A in the WTs. To further investigate the flexibility of the C-terminal lysines in the multiple mutants, the distance between each lysine (or alanine) was examined (Fig. S15). In the double mutant, both H2AK125 and H2AK129 are the lysines that travel the closest to the E2 active site (Fig. S15a), while H2AK127 is farther by ∼5 Å. In the triple mutant systems, in which H3 has no impact on the dynamics of the protein, all H2A C-terminal residues appear within the same proximity to the E2 active site, with H2AK119 being the closest lysine (Fig. S15b). The results of the multiple mutants possibly suggest that the interaction of H3 with Ub orchestrates conformational changes of the complex to favor the ubiquitination of the last three lysines on the C-terminal tail. This proposition is further supported by the presence of both the H3-Ub and H3-H2A contact in the double mutant system (Table S3), while the triple mutant system saw no contact with H3 for the duration of its simulation time. Moreover, the distance analysis of the double mutant (Fig. S15a) suggests that out of the 3-terminal lysine, H2AK125 and H2AK129 show a more probable proximity to the E2 active site over H2AK127. This further supports the results seen in the WT trials that when the H3-Ub interaction occurs, H2AK125 appears to be favored for ubiquitination, followed by H2AK129. In summary, based on the mutant results, we propose that the H3-Ub interaction may be crucial to the preferential ubiquitination of one of the last three lysines on the H2A C-terminal tail, as reported by experimental results.^[7,10,12,17,34]^

## Conclusion

The ubiquitination of the H2A C-terminal by the BRCA1/BARD1-UbcH5c (E3-E2) complex remains an interest of study due to the role of the process in cancer incidence. In the current study, we sought to provide insight into the role of human Ub on the structural dynamics of the E3-E2 system, as well as the contribution of H3 for selective ubiquitination of H2A lysines. The tilt angles and distance analysis suggested that Ub confines the BRCA1/BARD1 and orients UbcH5c/Ub at the feasible distance for ubiquitination. PCA and distance analysis reveal asymmetric movements between the E2 and Ub, thus providing further insight into the relationship between the two components in the system’s dynamics. Although results depict that H2AK119 is generally the closest lysine to the E2 active site, we discovered that it relies on the interactions established by H3. For instance, WT simulations reveal that if an interaction occurs between the N-terminus of H3 and Ub, H2AK125 is favored for the closest proximity to the E2 active site; however, when the H3-Ub interaction is absent, H2AK119 remains favored. Experimental research reports that the BRCA1/BARD1-UbcH5c complex preferentially ubiquitinates the last three lysines (H2AK125, H2AK127 and H2AK129).^[7,10,12]^ In our study, we have found that while these lysines approach the E2 active site much less than H2AK119, their role in the dynamics of the system is of great importance. Furthermore, the double mutant, H2AK118/119A, mutations have minimal effect on the contact frequency between E2 and H2A, while the triple mutant system of H2AK125/127/129A exhibits significantly diminished contact frequencies. Furthermore, the presence of the H3-Ub interaction allows for Ub to be brought closer to the NCP surface and allows E2 to move away from the NCP surface. When E2 moves away from the NCP surface, the last three lysine are able to move closer to the E2 active site. The flexibility of the C-terminal tail of H2A when E2 is elevated is consistent with the dynamics reported in the Ub-free system.^[11]^ However, in the absence of the H3-Ub interaction, the flexibility of the C-terminal tail is diminished, and thus it becomes more taxing for the last three lysine to approach the active site of E2. Since MD simulations cannot model the breaking and forming of bonds, the transfer of ubiquitin to H2A lysines could not be modeled in this study. For future studies, using QM/MM methods would provide further information into the energetics of the transfer of Ub to one of the H2A C-terminal lysines. Additionally, further investigation into the observed H3-Ub interaction would be beneficial for further insight into the possible role of H3 in BRCA1/BARD1-UbcH5c lysine preference.

## Acknowledgments

The authors acknowledge the Office of Information Technology Cyberinfrastructure Research Computing at the University of Texas at Dallas for providing HPC resources to conduct simulations and analyses for this study. Additionally, the authors acknowledge Dr. Horn at Friedrich-Alexander Technology Universität Erlangen-Nürnberg, who developed and provided the isopeptide bond parameters used in this work. A startup fund from the University of Texas at Dallas was utilized to fund this work.

## Author’s Contribution

H.T. designed the project; A.R.G. and T.S. prepared the model; A.R.G. performed the computational studies and analysis; H.T., T.S., and A.R.G. analyzed the data and wrote the manuscript together.

## Data Availability

The input files, parameters, trajectories, and other data will be provided upon request.

