## Supplementary Figures for "Histone H3 Orchestrates the Ubiquitination of Nucleosomal H2A by BRCA1/BARD1-UbcH5c Complex"

University of Texas at Dallas

800 W. Campbell Rd., Richardson, TX

**Additional Methods**

I. **Model Preparation**

AMBER force fields ff14SB^[1]^, DNA.OL15^[2]^, and GAFF2^[3]^ were used for proteins, nucleic acids and organic molecules, respectively. Furthermore, the four tetrahedrally coordinated zincs at the base of BRCA1/BARD1 were modeled using ZAFF^[4]^ to properly model the bonds between each zine and its respective cysteine and histidine residues. Upon bonding Ub, the system was then explicitly solvated with TIP3P water^[5]^ and was explicitly neutralized with 150mM of NaCl. Additionally, double and triple mutant systems were prepared to investigate the ubiquitination selectivity and electrostatic interactions of the H2A tail. All mutants were prepared in the same fashion as the wild type (WT) system, utilizing AMBER’s Leap program. All systems, including the WT, under AA-MD simulations.

II. **Simulation Details**

AA-MD simulations were conducted for all WT and mutated systems of the BRCA1/BARD1-UbcH5c NCP with bound Ub structure. Three trials were conducted for the WT system, all starting with the same structure but different initial velocities in order to provide randomization and reliable results. The double mutant system (K118A and K119A) and the triple mutant system (K125A, K127A, K129A) ran for only one trial. Each WT simulation ran for a total of 1 μs of production, whereas mutant systems ran for a total of 500 ns of production. Before production, all systems underwent minimization, heating, and equilibration, and the methodology of these phases remained the same as that used for the previous system.

Analysis including root mean square deviation (RMSD) of the protein backbone, side chain root mean square fluctuations (RMSF), hydrogen bonding, tilt angle, PCA, and all distance analysis were performed using AMBER20’s CPPTRAJ module. Furthermore, the normalization of tilt angle and distance results were also conducted using the CPPTRAJ module. The tilt angle was determined in the same fashion as in the previous publication. Two vectors were defined by an alpha helix of H2B on the NCP surface, and an alpha helix of BRCA1 or BARD1, and the dot product of the two vectors was taken. Salt bridge analysis was conducted using the Salt Bridges Plugin^[6]^ extension in VMD.^[7]^ The extension indicates salt bridges based on a 3.2Å cutoff between oxygen and nitrogen atoms on differing residues. Finally, contact frequency analysis between E2 and the H2A C-terminal tail was performed through VMD^20^ using the in-house TCL code.

**Table S1:** Residue Ranges for Protein Components

| **Component** | **Residue Range** |
| --- | --- |
| BRCA1-UbcH5c | 1-260 |
| BARD1 | 261-359 |
| H2A | 360-629 |
| H2B | 630-883 |
| H3 | 884-1157 |
| H4 | 1158-1365 |
| DNA | 1366-1659 |
| Zinc | 1660-1663 |
| Ub | 1664-1740 |

**Table S2:** Total Simulation Times

| **System** | **Trial** | **Production Time (ns)** |
| --- | --- | --- |
| WT | 1 | 1,000 |
| WT | 2 | 1,000 |
| WT | 3 | 1,000 |
| K118/119A (Double Mutant) | 1 | 500 |
| K125/127/129A (Triple Mutant) | 1 | 500 |

**Table S3:** H3 Interaction Occurrence Across All Systems.^a^

| System | Trial | H3-Ub | H3-NCP |
| --- | --- | --- | --- |
| WT | 1 | 81.89% | 10.71% |
|  | 2 | 0.00% | 75.72% |
|  | 3 | 0.25% | 3.74% |
| K118/119A | 1 | 58.93% | 49.13% |
| K125/127/129A | 1 | 0.00% | 0.00% |

^a^The occurrence is computed as the average contact frequency between the N-terminal tail of H3 and with residues 57 through 64 on Ub (H3-Ub interaction) and the 5 C-terminal lysines on H2A (H3-NCP interaction).

**Table S4:** Modes of PCA for WT Systems^b^

| Mode |  | WT 1 | WT 2 | WT 3 | Sum of % Contribution | Mode Description |
| --- | --- | --- | --- | --- | --- | --- |
| A | 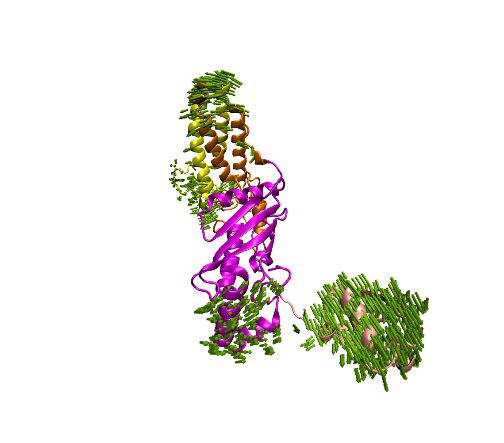 | - | PC 1  56.33% | PC 1  36.05% | 92.38% | BRCA1/BARD1: Tilt  E2/Ub: Seesaw Motion |
| B | 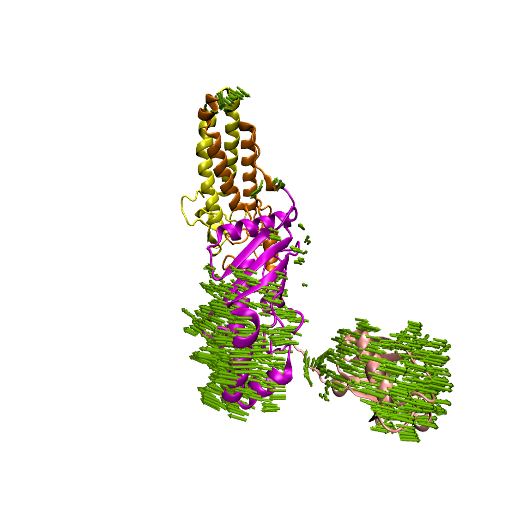 | PC 2  20.03% | PC 2  16.38% | - | 36.41% | BRCA1/BARD1: Prominent tilt  E2/Ub: Rotation |
| C | 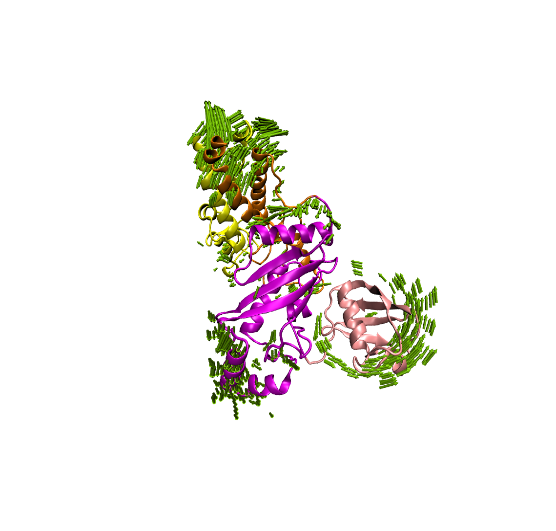 | PC 1  27.71% | - | - | 27.71% | BRCA1/BARD1: tilt  E2/Ub:Hinging of E2 with Sliding of Ub |

^b^Figures of PC modes show movements of 2 Å or greater.

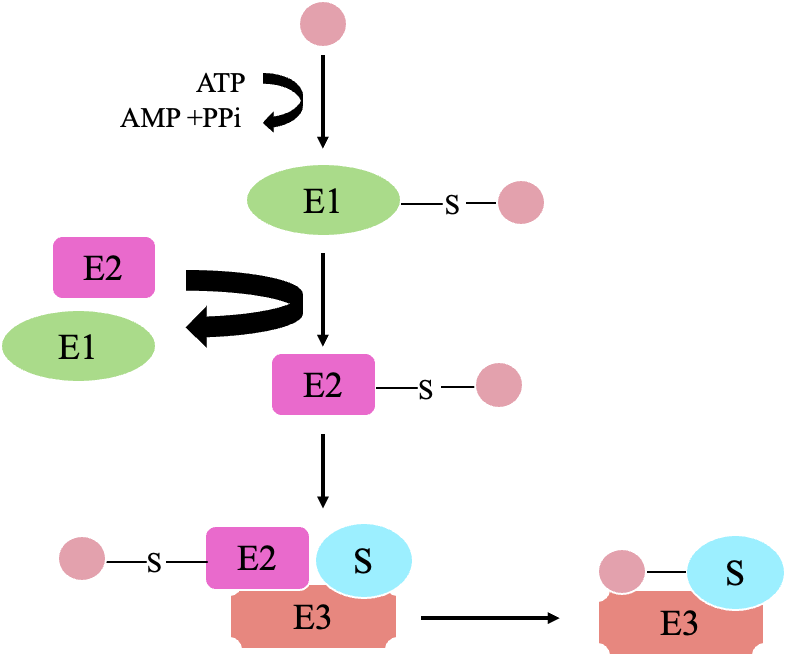

**Fig. S1:** Ubiquitination Process Scheme. Ubiquitin (pink) forms thioester bond with the ubiquitin activating enzyme (E1, green) through the ATP dependent first step known as activation. In the second step, conjugation, ubiquitin is transferred from E1 to the ubiquitin conjugating enzyme (E2, magenta) where ubiquitin forms a thioester bond with E2’s active site. In the final step, ligation, the ubiquitin ligase (E3, salmon) catalyzes the transfer of ubiquitin from E2 to the target substrate (cyan) where the C terminal glycine of ubiquitin forms an isopeptide bond with a lysine on the substrate.

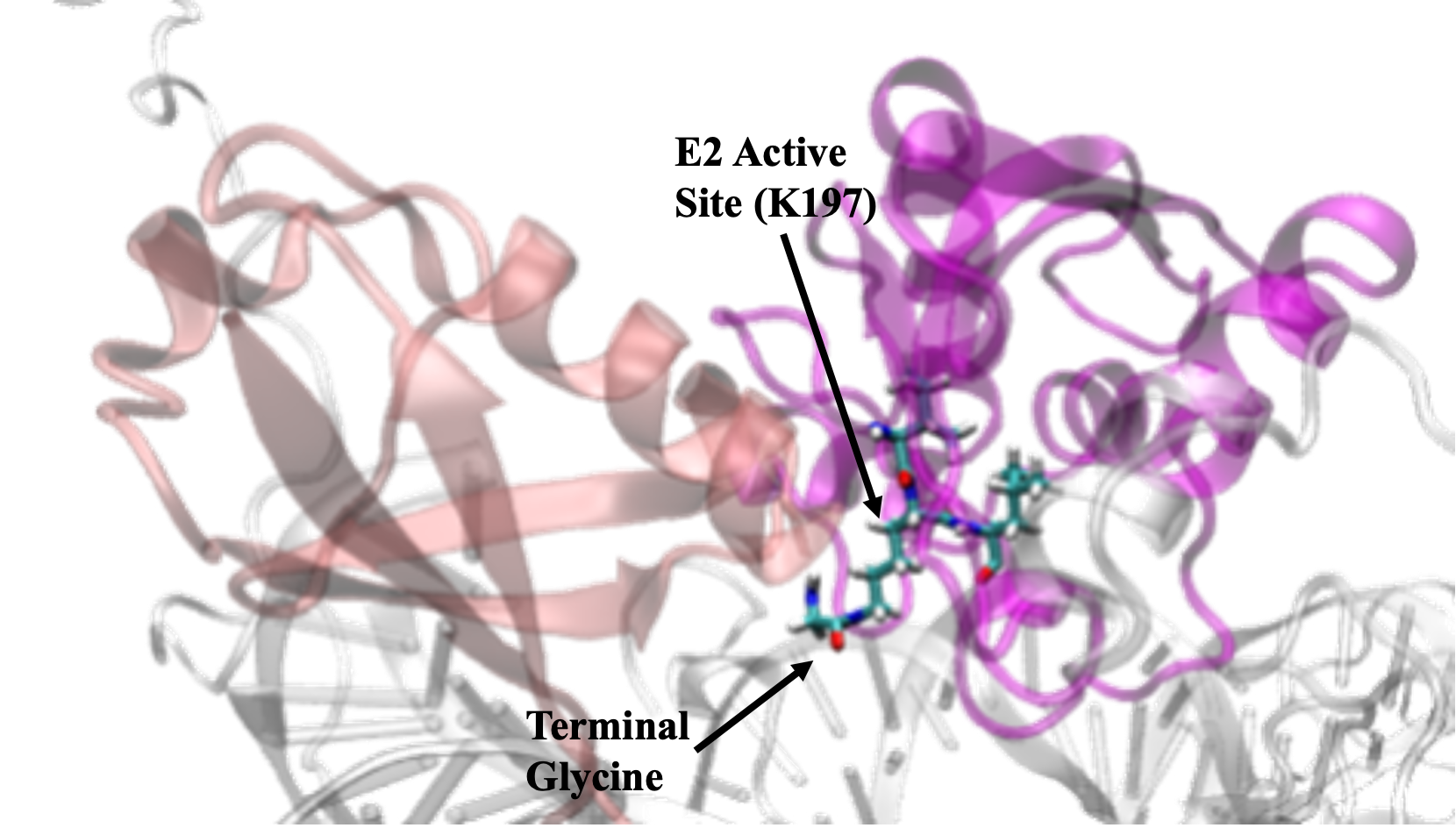

**Fig. S2:** Isopeptide Bond. The terminal glycine of Ub (G76) (pink) was bonded to the E2 active site (K197) (magenta) via an isopeptide bond.

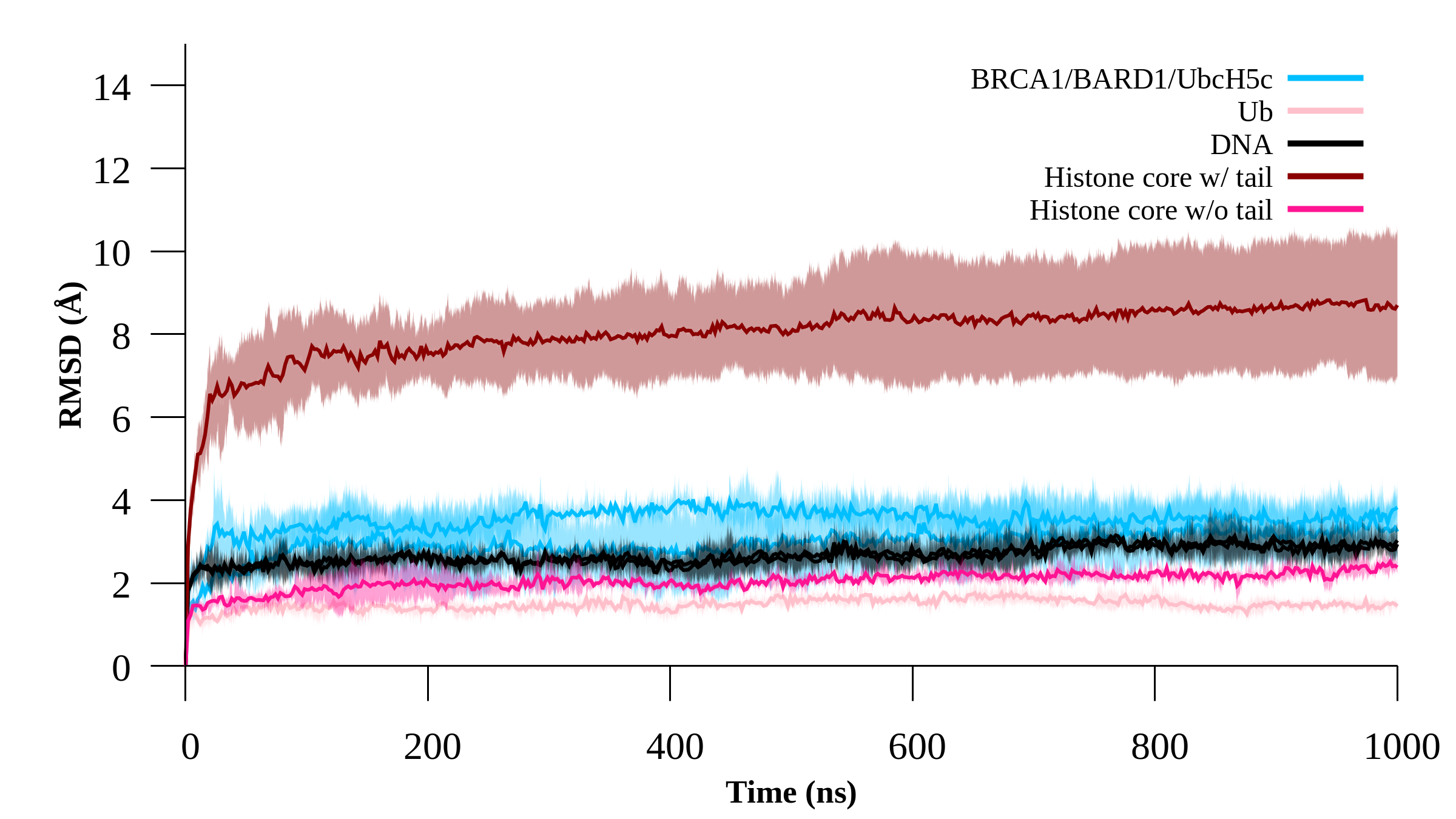

**Fig. S3:** RMSD of BRCA1/BARD1/UbcH5c (cyan), Ub (pink), DNA (black), and histones with (magenta) and without (brown) tails for WT simulations. All shading indicates the standard deviation of the data set.

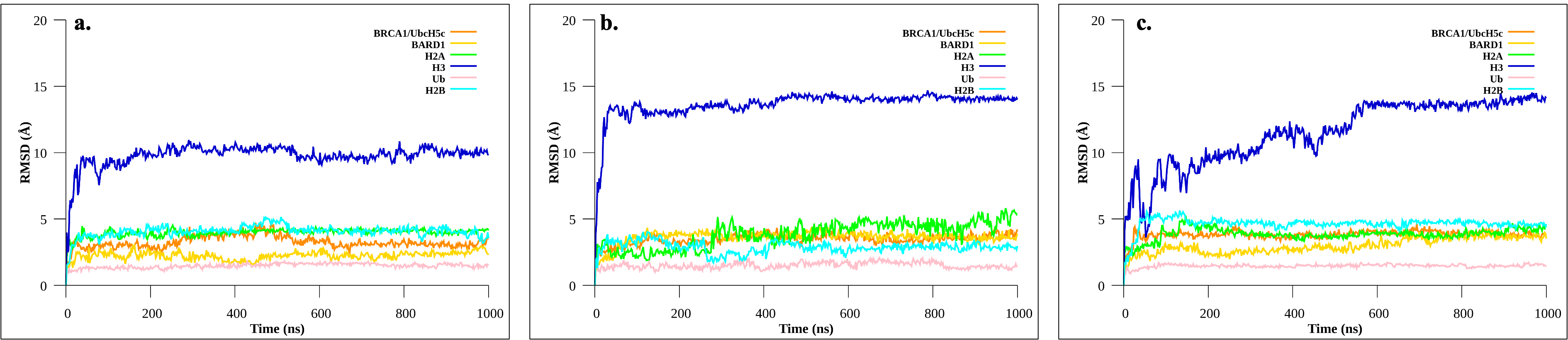

**Fig. S4:** RMSD for Structure of Interest. (a) RMSD for WT trial 1 of BRCA1/UbcH5c (orange), BARD1 (yellow), Ub (pink), H2A with disorder tail (green) and H3 with disordered tail (blue). (b) RMSD of same regions for WT 2. (c) RMSD of same regions for WT 3.

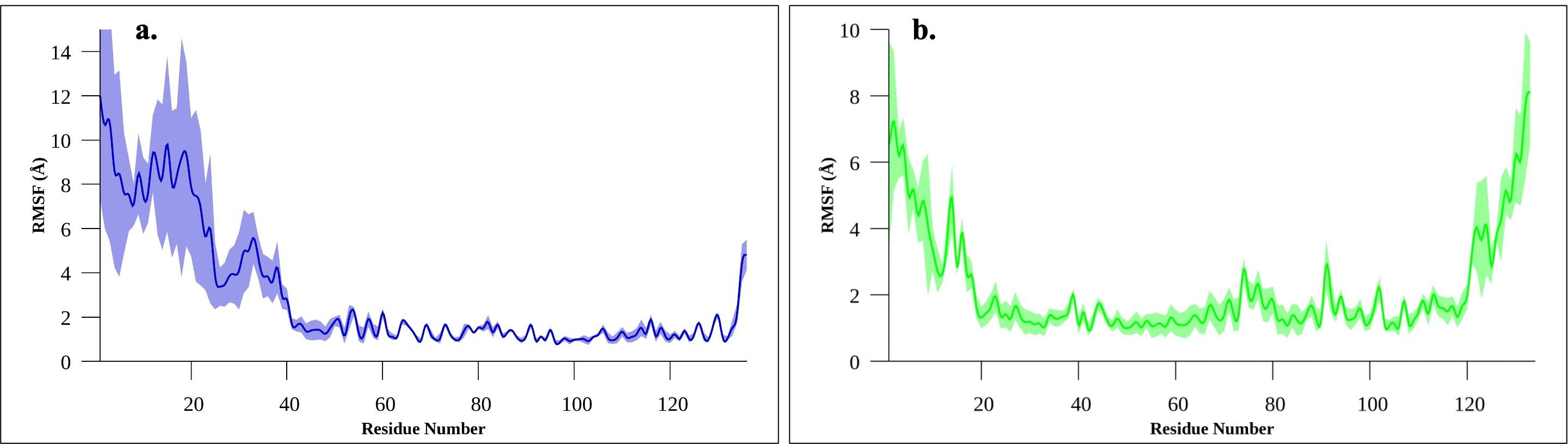

**Fig.S5:** RMSFs for Crucial Histones. (a) RMSF of H3. (b) RMSF of H2A.

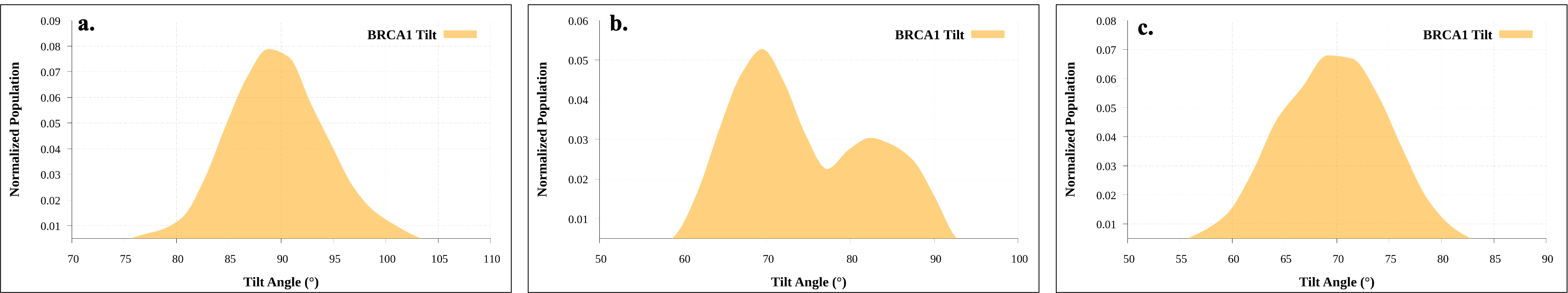

**Fig. S6:** BRCA1 Tilt Angles for WT Trials. (a) BRCA1 tilt angle for WT 1. (b) BRCA1 tilt angle for WT 2. (c) BRCA1 tilt angle for WT 3.

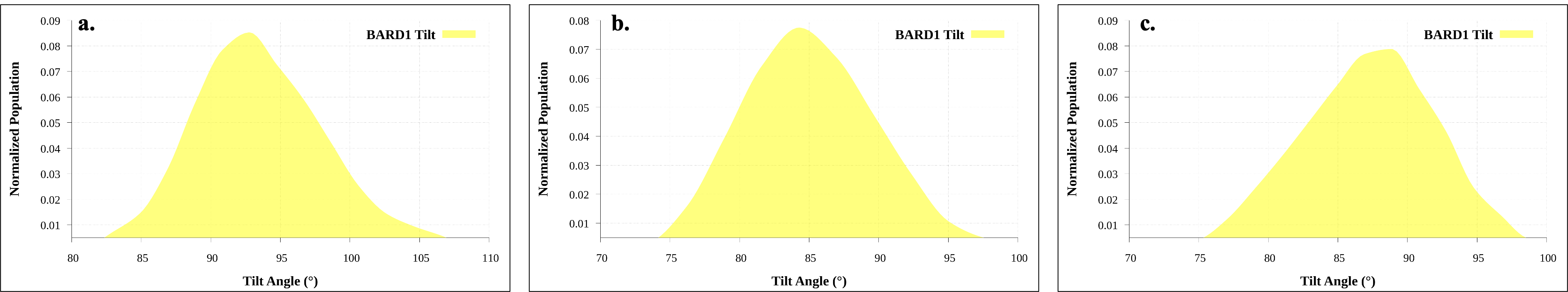

**Fig. S7:** BARD1 Tilt Angles for WT Trials. (a) BARD1 tilt angle for WT 1. (b) BARD1 tilt angle for WT 2. (c) BARD1 tilt angle for WT 3.

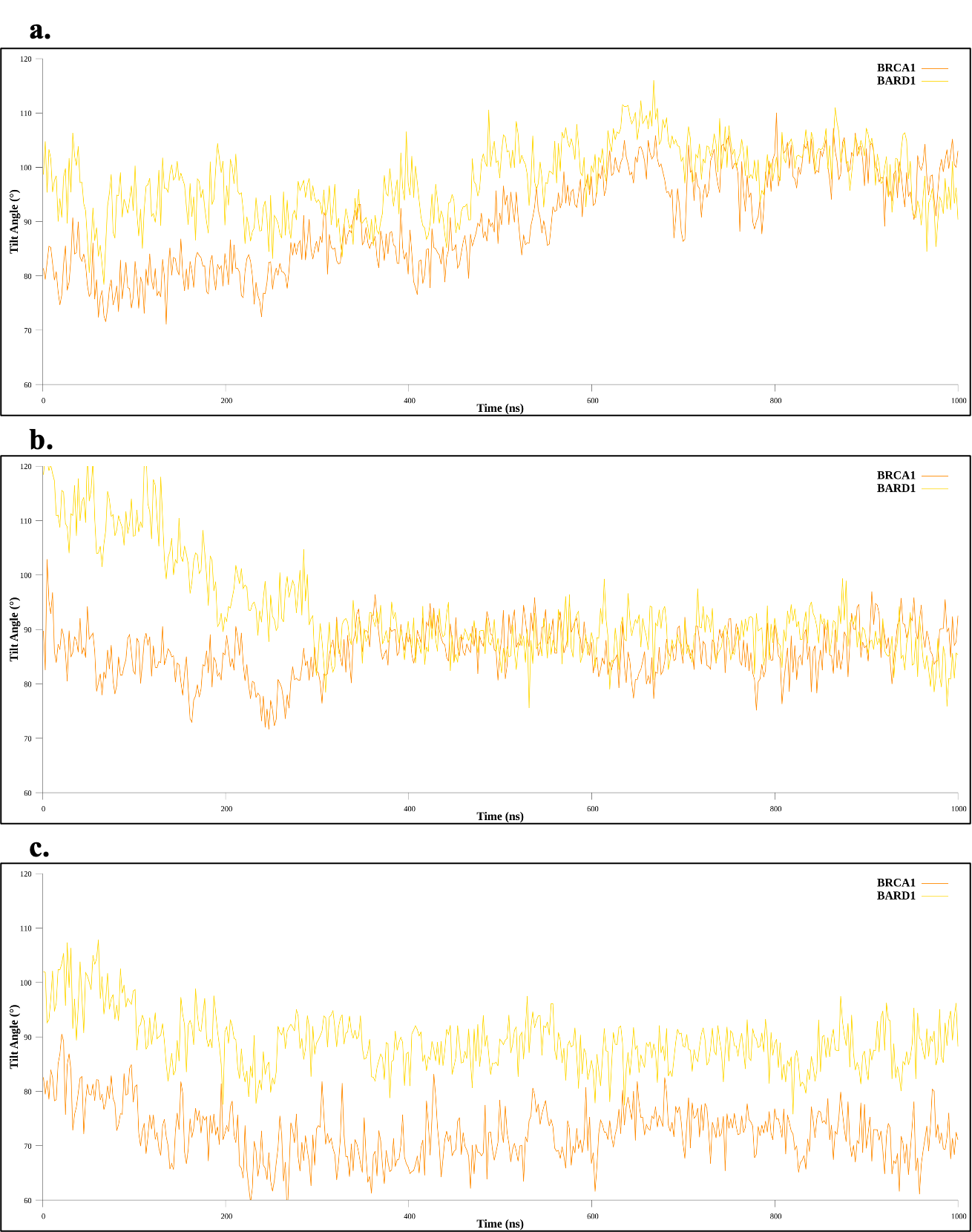

**Fig. S8:** Tilt Angle as a Function of Time. Tilt angle of both BRCA1 (orange) and BARD1 (yellow) over the 1000ns of production for (a) WT trial 1, (b) WT trial 2 and (c) WT trial 3.

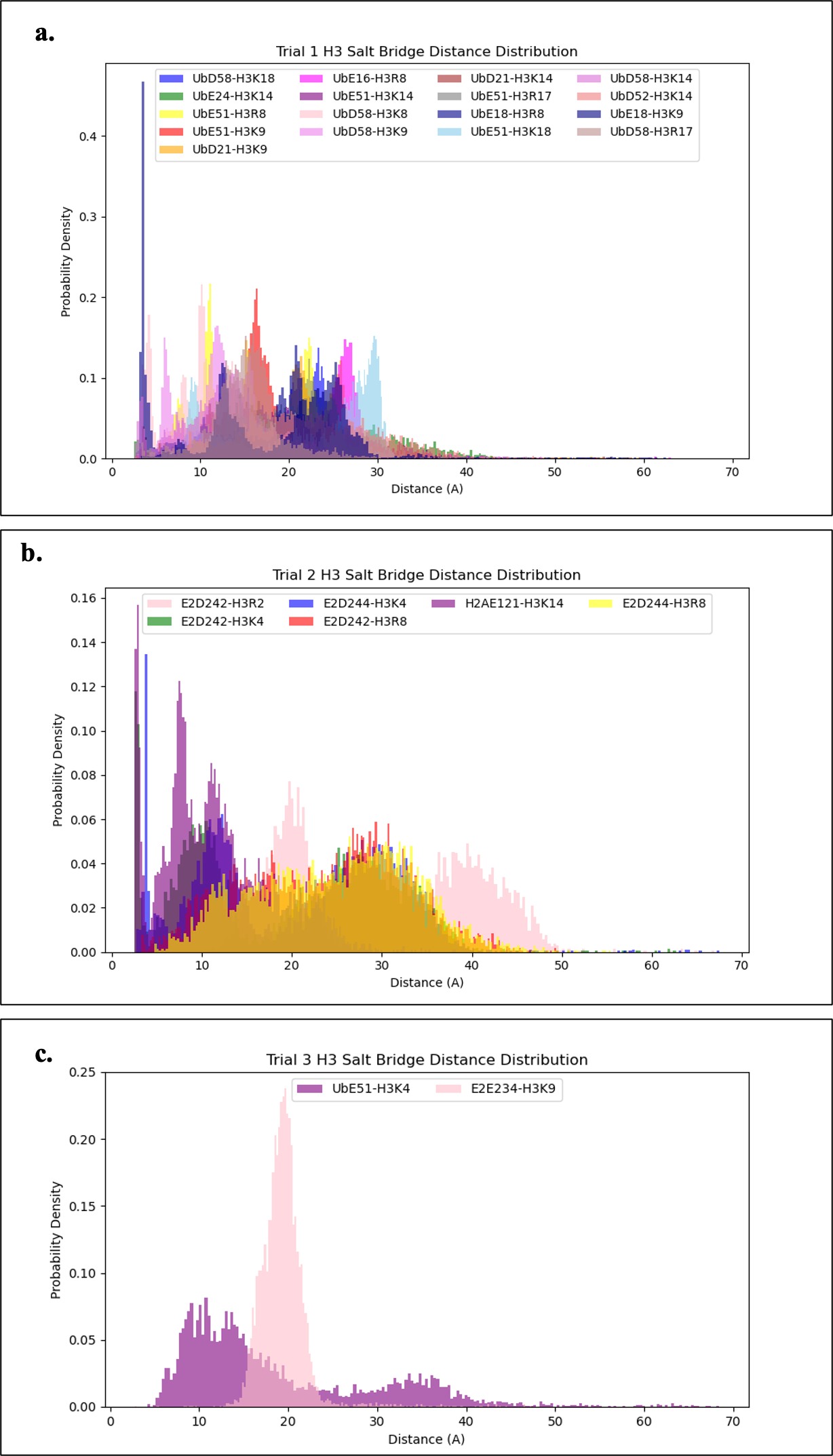

**Fig. S9:** H3 N-Terminal Tail Salt Bridges. The distribution curves show the salt bridges formed between the first 20 residues of H3 N-terminal tail with different residues on the protein in WT trials (a) 1, (b) 2, and (c) 3. The naming convention in the legend of each plot follows “portion of protein, residue on protein”

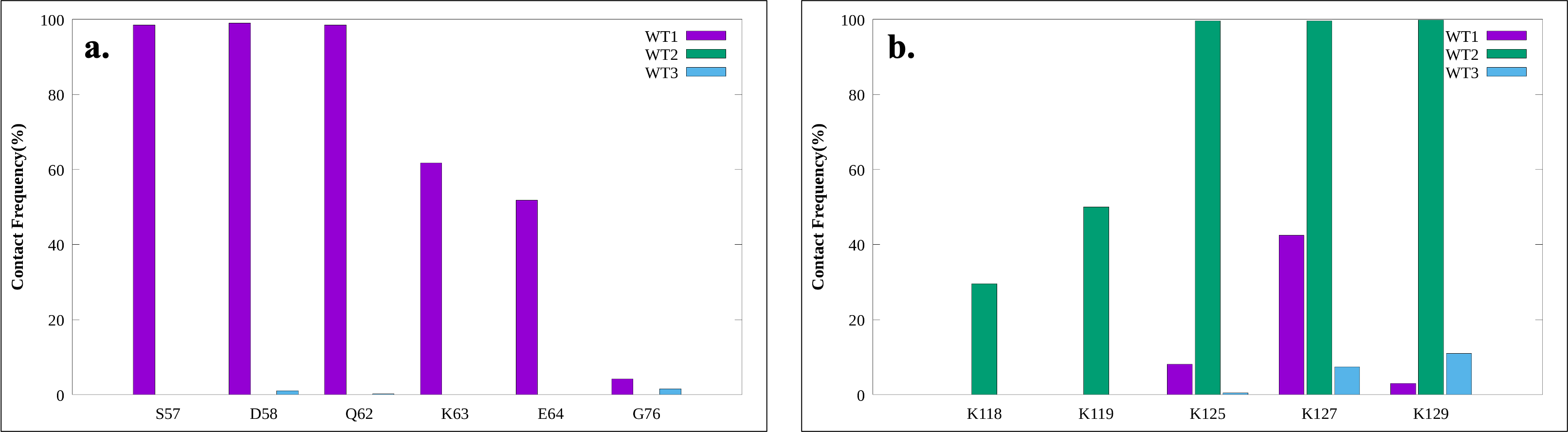

**Fig. S10:** H3 N-Terminal Tail Contact Frequency. H3 contact across all WT trials with (a) Ub and (b) H2A.

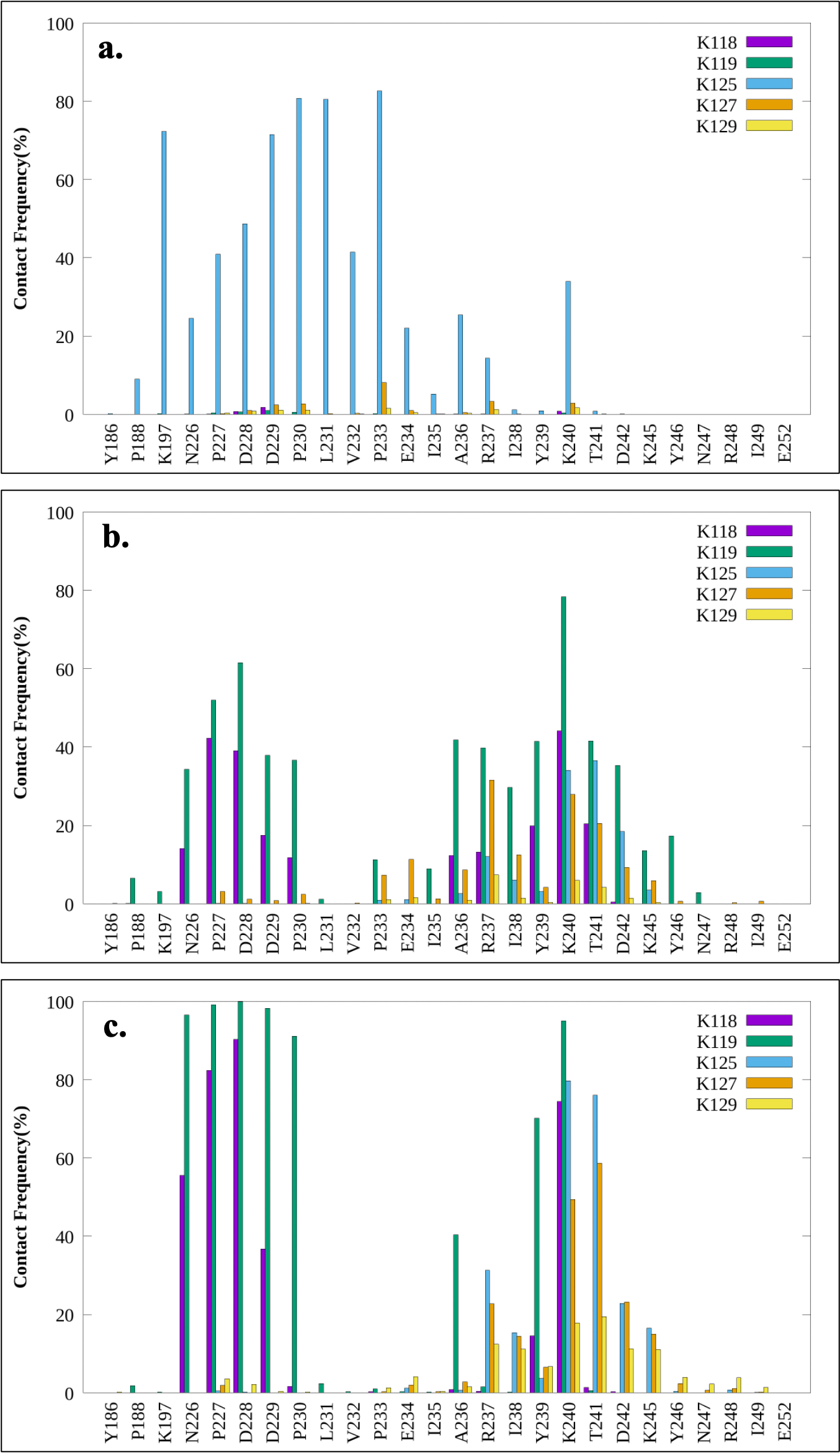

**Fig. S11:** Contact Frequency of WT Systems. Data shows the contact frequency for (a) WT trial 1 (b) WT trial 2 and (c) WT trial 3 between H2A C-terminal lysines and residues on BRCA1/UbcH5c

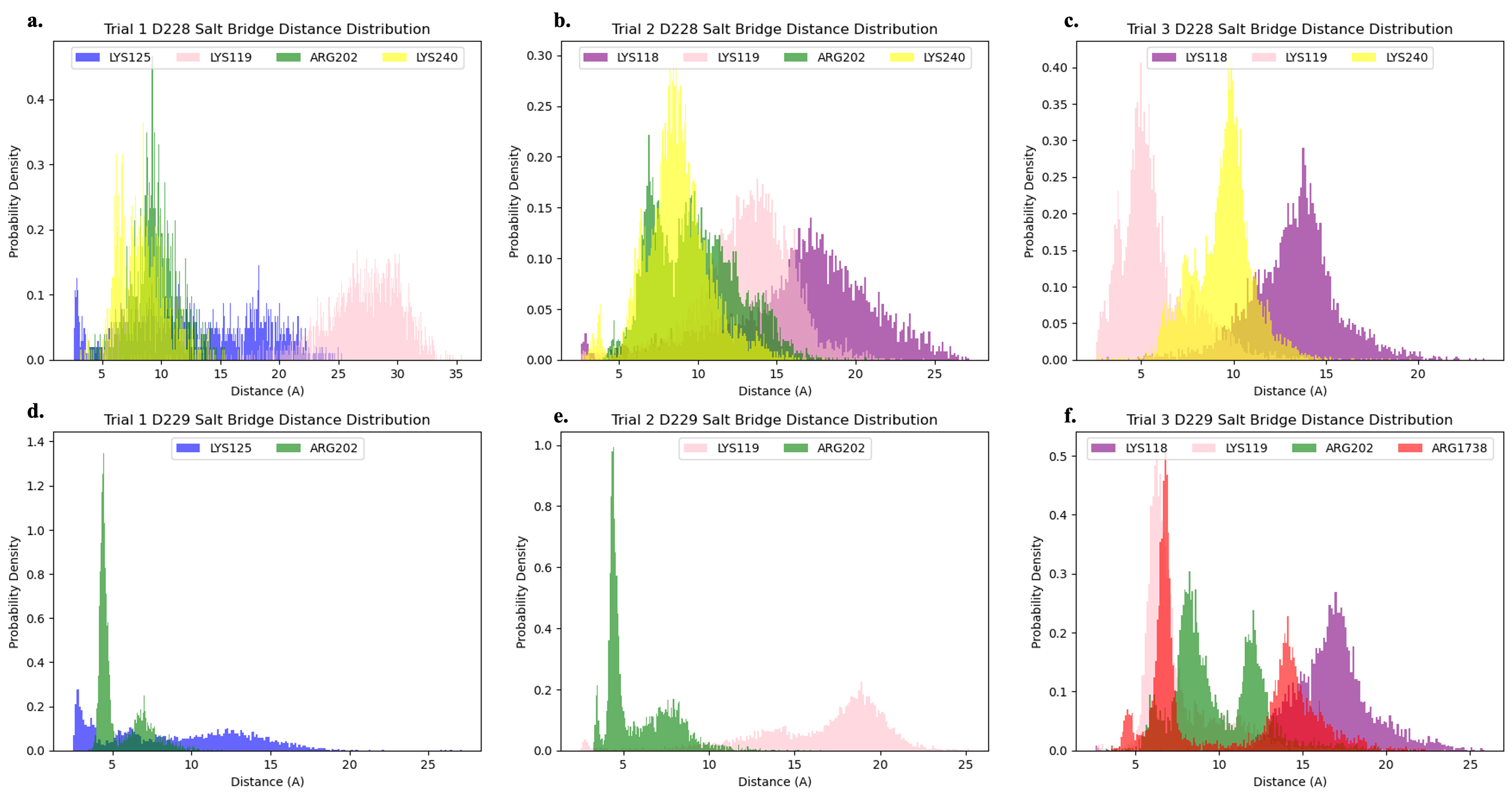

**Fig. S12:** Salt Bridge Analysis For D228 and D229. (a) Salt bridges formed with D228 in WT trial 1. (b) Salt bridges formed with D228 in WT trial 2. (c) Salt bridges formed with D228 in WT trial 3. (d) Salt bridges formed with D229 in WT trial 1. (e) Salt bridges formed with D229 in WT trial 2. (f) Salt bridges formed with D229 in WT trial 3.

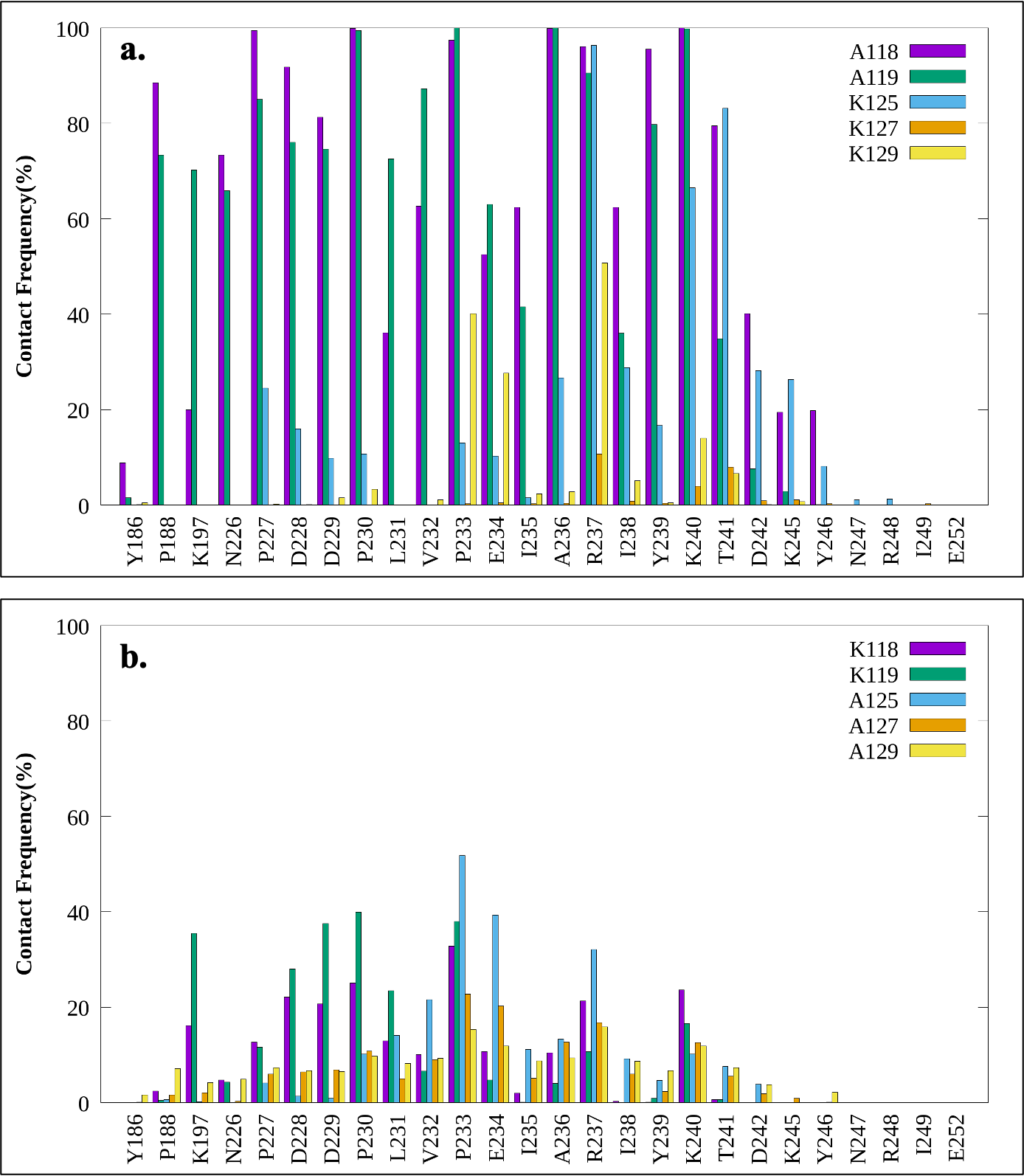

**Fig. S13:** Multiple Mutant E2-H2A Contact Frequency. (a) Contact frequency for the double mutant system (K118A and K119A) with E2. (b) Contact frequency for the triple mutant system (K125A, K127A and K129A) with E2.

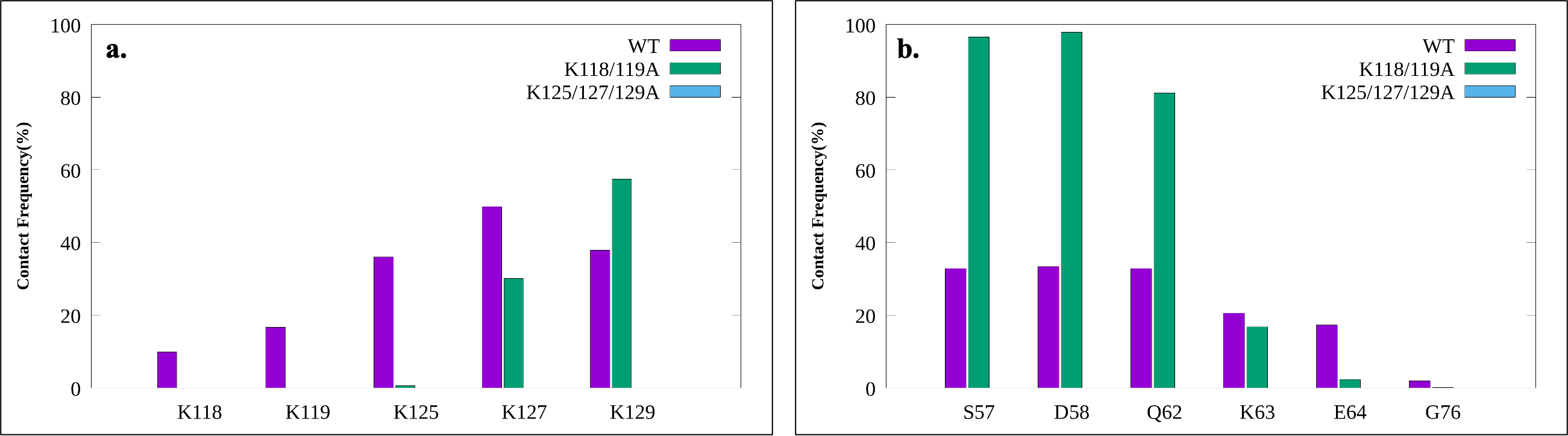

**Fig. S14:** Contact Frequency with H3 Tail. (a) The contact frequency between the H3 disordered region on H3 in the WT (purple), double mutant (green), and triple mutant (blue) with H2A C-terminal lysines. (b) The contact frequency between H3 and ubiquitin residues with the largest RMSFs for the aforementioned 3 systems.

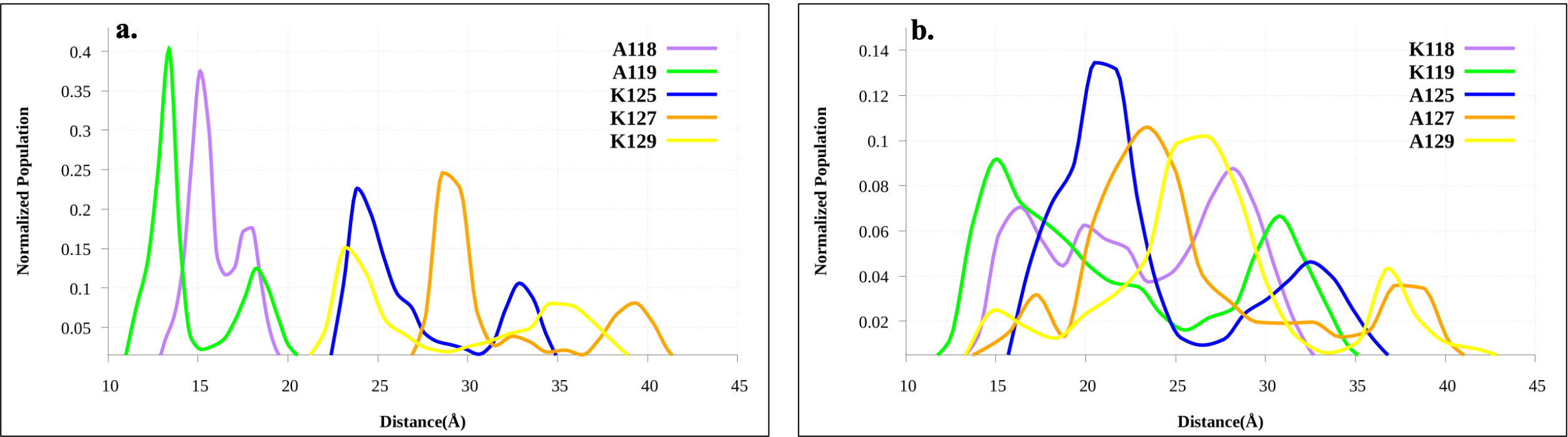

**Fig. S15:** H2A Lysine to E2 Active Site Distance of Multiple Mutants. (a) Lysine to E2 active site distance for the double mutant. (b) Lysine to E2 active site distance for triple mutant.
